# Modeling the efficacy of CRISPR gene drive for schistosomiasis control

**DOI:** 10.1101/2021.10.29.466423

**Authors:** Richard E. Grewelle, Javier Perez-Saez, Josh Tycko, Erica K.O. Namigai, Chloe G. Rickards, Giulio A. De Leo

## Abstract

CRISPR gene drives could revolutionize the control of infectious diseases by accelerating the spread of engineered traits that limit parasite transmission in wild populations. While much effort has been spent developing gene drives in mosquitoes, gene drive technology in molluscs has received little attention despite the role of freshwater snails as obligate, intermediate hosts of parasitic flukes causing schistosomiasis – a disease of poverty affecting more than 200 million people worldwide. A successful drive in snails must overcome self-fertilization, which prevents a drive’s spread. Simultaneous hermaphroditism is a feature of snails – distinct from gene drive model organisms – and is not yet incorporated in gene drive models of disease control. Here we developed a novel population genetic model accounting for snails’ sexual and asexual reproduction, susceptibility to parasite infection regulated by multiple alleles, fitness differences between genotypes, and a range of drive characteristics. We then integrated this model with an epidemiological model of schistosomiasis transmission and snail population dynamics. Simulations showed that gene drive establishment can be hindered by a variety of biological and ecological factors, including selfing. However, our model suggests that, under a range of conditions, gene drive mediated immunity in snails could maintain rapid disease reduction achieved by annual chemotherapy treatment of the human population, leading to long-term elimination. These results indicate that gene drives, in coordination with existing public health measures, may become a useful tool to reduce schistosomiasis burden in selected transmission settings with effective CRISPR construct design and close evaluation of the genetic and ecological landscape.

## Introduction

Gene drive technology is rapidly expanding since the discovery of CRISPR-Cas9 [1–3]. Its potential uses include controlling diseases, invasive species, and pests by spreading targeted genes through a population faster than traditional Mendelian inheritance allows [4]. For example, there are currently large efforts to harness genetic technology targeting mosquito species that are vectors of malaria and other vector-borne diseases [5–9]. Similar efforts could be on the horizon for schistosomiasis, a debilitating disease of poverty caused by blood flukes of the genus Schistosoma [10].

The battle to eliminate schistosomiasis has been waged for more than a century, and despite local successes, the disease remains widespread [11]. Globally over 200 million individuals are actively infected. With 800 million people at risk of infection, schistosomiasis is second only to malaria in the breadth of its health and economic impact as an infectious tropical disease [12, 13]. The disease manifests as a complex suite of symptoms stemming primarily from the inflammatory processes the body mounts in response to the schistosome eggs that embed in tissue [14]. Abdominal pain, release of blood in urine or stool, fever, enlargement of liver or spleen, and accumulation of fluid in the peritoneal cavity are acute symptoms, while fibrosis and lesions of vital organs, infertility, and several forms of cancer are lasting consequences of infection [15, 16].

Transmission of schistosomes to intermediate, obligate snail hosts occurs when eggs shed in urine or feces from infected people contact freshwater and emerge as free-swimming miracidia. Once established within the snail, the parasite reproduces asexually and cercariae are released 3-5 weeks after the onset of infection. In this stage, the parasites castrate the freshwater snails, severely reducing reproduction [17]. Released cercariae can penetrate the skin of humans in contact with infested water bodies and cause infection (Fig 1) [18].

**Figure 1:**
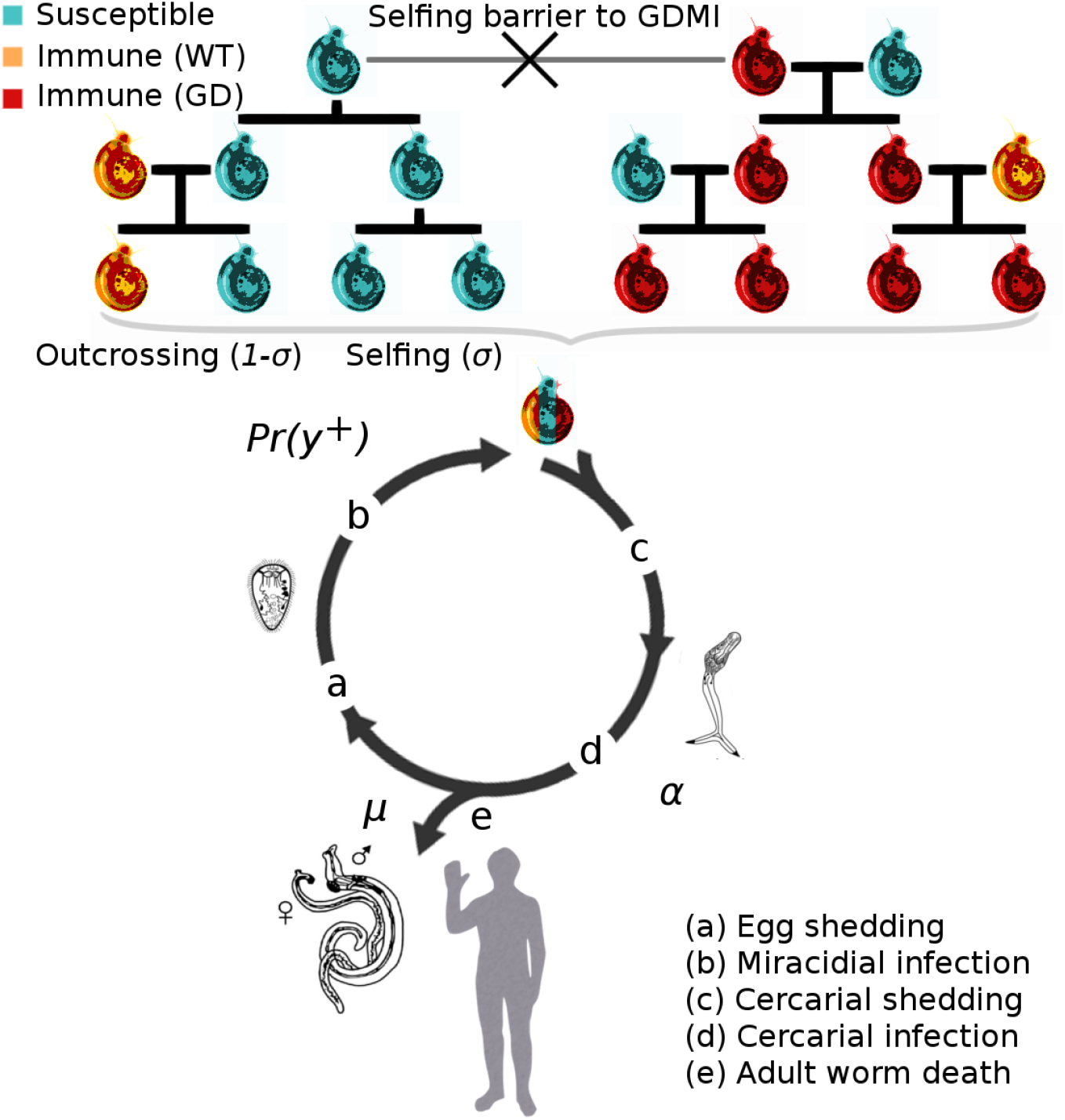
Conceptual diagram of the integrated epidemiological and population genetic model describing the evolution of immunity to schistosome infection in the snail population. (a) High worm burden in the human population increases the force of infection on the snail population, which positively selects for immune snail genotypes. (b) Miricidial infection of susceptible snails is density-dependent. (c) Evolution of immunity in the snail population reduces cercarial transmission to humans, thereby regulating parasite densities at an endemic equilibrium. GDMI is inherited more rapidly than natural immunity only when outcrossing occurs. (d) Infection of humans is proportional to cercarial output, and a negative binomial distribution of adult worms in the human population influences mating success and egg production. Immunity in the human population is assumed constant due to accumulated evidence that immunity acquisition occurs over decades and likely varies little relative to immunity in the snail population in a 10 year window in endemic conditions. (e) Mortality of adult worms occurs via constant natural mortality and MDA treatment. Three snail genotypes are modeled: susceptible to infection, innately immune (wild type), and gene drive mediated immune. Iteroparitive reproduction and mortality of these genotypes is modeled with explicit fecundity and viability components of fitness (see SI).

Rapid advancements in genomics for the intermediate snail host species provides a mechanistic understanding of innate, genetically-based snail immunity to schistosome infection [19]. Genes responsible for immunity could be candidates for gene drive mediated spread through snail host populations. Promisingly, selection experiments reveal rapid evolution of immune phenotypes, demonstrating high immunity can be achieved under laboratory conditions within a few snail generations [20, 21] (SI Fig. S3). Overall, there is good reason to expect that a CRISPR gene drive designed to provide greater immunity in the snail population could soon be developed. However, whether such a gene drive could provide the sustained reduction in transmission necessary to eliminate schistosomiasis in realistic settings laden with barriers to the spread of a drive remains unknown.

Previous theoretical work using classical population genetic models has explored how fitness, homing efficiency, selfing, resistance allele formation, gene flow, and other forms of population structure influence invasion success and peak frequency of a drive in general contexts [22–24]. Stochastic Moran models or discrete deterministic models with non-overlapping generations do not incorporate population dynamics on which the tempo of evolution is highly dependent. Snail populations are iteroparous, reproducing several times within a lifetime, and exhibit density dependent recruitment. This form of reproduction is not modeled in the simplified evolutionary models developed for gene drives to date. Accuracy of gene drive models hinges on realistic assumptions of the target population.

The success of gene drive mediated immunity (GDMI) in natural snail populations is determined by features intrinsic to the design of the drive construct and its deployment – homing efficiency, fitness cost of the payload, evolution of resistance to the drive, and number of releases – and by extrinsic properties of the environment in which GDMI is deployed, such as the size of the focal snail population, transmission rates, and gene flow and standing genetic variation for immunity in snail populations. Importantly, all snail species that serve as intermediate hosts to schistosomes, except for *Oncomelania spp.*, are simultaneous hermaphrodites capable of self-fertilization (selfing). In contrast to mosquito and fruit fly models for which gene drives have been designed, selfing snail species may be incapable of propagating a drive construct. Gene drive relies on an encoded endonuclease, such as Cas9, which introduces a double strand break in the homologous chromosome that is repaired using the gene drive allele as a template, thereby copying the gene drive allele to the homologous chromosome [4, 5]. Sexual reproduction (outcrossing) is necessary for gene drive to spread a target allele in a population through pairing and reassortment of gene drive and wild type alleles, facilitating gene conversion. Because the propensity to self-fertilize varies by species and environmental conditions [25], it is imperative to understand how selfing interacts with the variety of intrinsic and extrinsic factors that may influence the establishment of GDMI in natural snail populations.

The impact of GDMI is determined by public health outcomes and not by establishment alone. Local success in schistosomiasis reduction can be achieved through sustained non-pharmaceutical intervention, but such approaches are often resource intensive (e.g. sanitation) or cause collateral damage to the environment (e.g. molluscicides) [26, 27]. Praziquantel (PZQ) emerged in the 1980s as the drug of choice for mass drug administration (MDA) [28, 29], and while cheap and effective in removing mature parasites from infected people and temporarily reducing morbidity, PZQ does not prevent reinfection, and extensive MDA campaigns have been unable to locally eliminate the disease in high transmission regions [30, 31]. For this reason, in recent years there has been a renewed interest in complementing MDA with environmental interventions aimed at targeting the environmental reservoirs of the disease [32–35]. GDMI has the potential to augment environmental interventions as a means toward cost-effective and sustainable schistosomiasis elimination, especially when paired with existing anthelmintic treatment of humans.

We investigate the role of selfing and its interaction with other factors influencing GDMI establishment to infer the challenges and opportunities for GDMI in a natural context. We hypothesized that a high selfing rate would incapacitate a gene drive, but a lower selfing rate could be compatible with a drive in certain conditions. To test these ideas we developed a biologically realistic mathematical model incorporating both genetic and environmental factors. This model is integrated in an epidemiological framework to evaluate the reduction of disease burden in humans with and without coincident MDA treatment. This study can be used as an informative first step for scientists, stakeholders, and policy makers looking to address the large human health crisis of schistosomiasis in conjunction with the principles for responsible use of gene drives proposed by the National Academies of Science, Engineering, and Medicine (NASEM) [36].

## Results

We developed a population genetic model that accounts separately for fecundity and viability components of fitness as well as for density dependent dynamics of the snail host population. We expand the wild type - gene drive, 2 allele model to separate the naturally occurring alleles into immune and susceptible types. The resulting six genotypes are formed from three alleles (susceptible, innately immune, gene drive mediated immune) and incorporated into a Markov model modified to include overlapping generations and population dynamics of susceptible and infected snails. Finally, we integrated the population genetic and ecological model with an epidemiological model to describe the dynamics of infection in the human population. Parameter values for the genetic model are derived from literature (SI Table S1) or otherwise explored in sensitivity analyses in the resulting figures, and under default conditions, simulated evolution recapitulates challenge experiments (SI Fig. S3). We examine the impact that the self-fertilization (selfing) rate of a focus population has on the establishment of gene drive in 10 years.

To represent the two species clusters, we depict results from both ends of the range of observed selfing rates among snail hosts. At high rates (σ = 0.8), self-fertilization undermines the gene drive and prevents establishment. However, GDMI is able to overcome low rates of self-fertilization (σ = 0.2) and establish at high frequencies (Fig. 2A). Self-fertilization is largely species dependent and may vary with local conditions with higher propensities to self-fertilize observed at low population densities [37]. To simulate this range of conditions, we perform a sensitivity analysis at the 10 year endpoint, so chosen as the likely window in which the efficacy of targeted treatments are evaluated in human populations. Especially for predominantly outcrossing species, reduced offspring viability is associated with self-fertilization [37]. We provide a confidence interval around the endpoint sensitivity analysis based on a range of inbreeding costs to fecundity. Gene drive success in 10 years is highly dependent on low selfing rate, though slower establishment is possible at moderate selfing rates. The inflection point near σ = 0.6 gives the value over which gene drive success is improbable in a 10 year window (Fig. 2B). These results indicate that for species with a lower rate of selfing, including *Oncomelania hupensis*, *Biomphalaria glabrata*, and *Bulinus globosus*, gene drive could establish rapidly in focus populations [38]. Conversely, for species like *Biomphalaria pfeifferi* or *Bulinus truncatus*, which have been observed to self-fertilize at rates higher than 0.6, gene drive will likely be ineffective in increasing immunity in the snail population [39].

**Figure 2:**
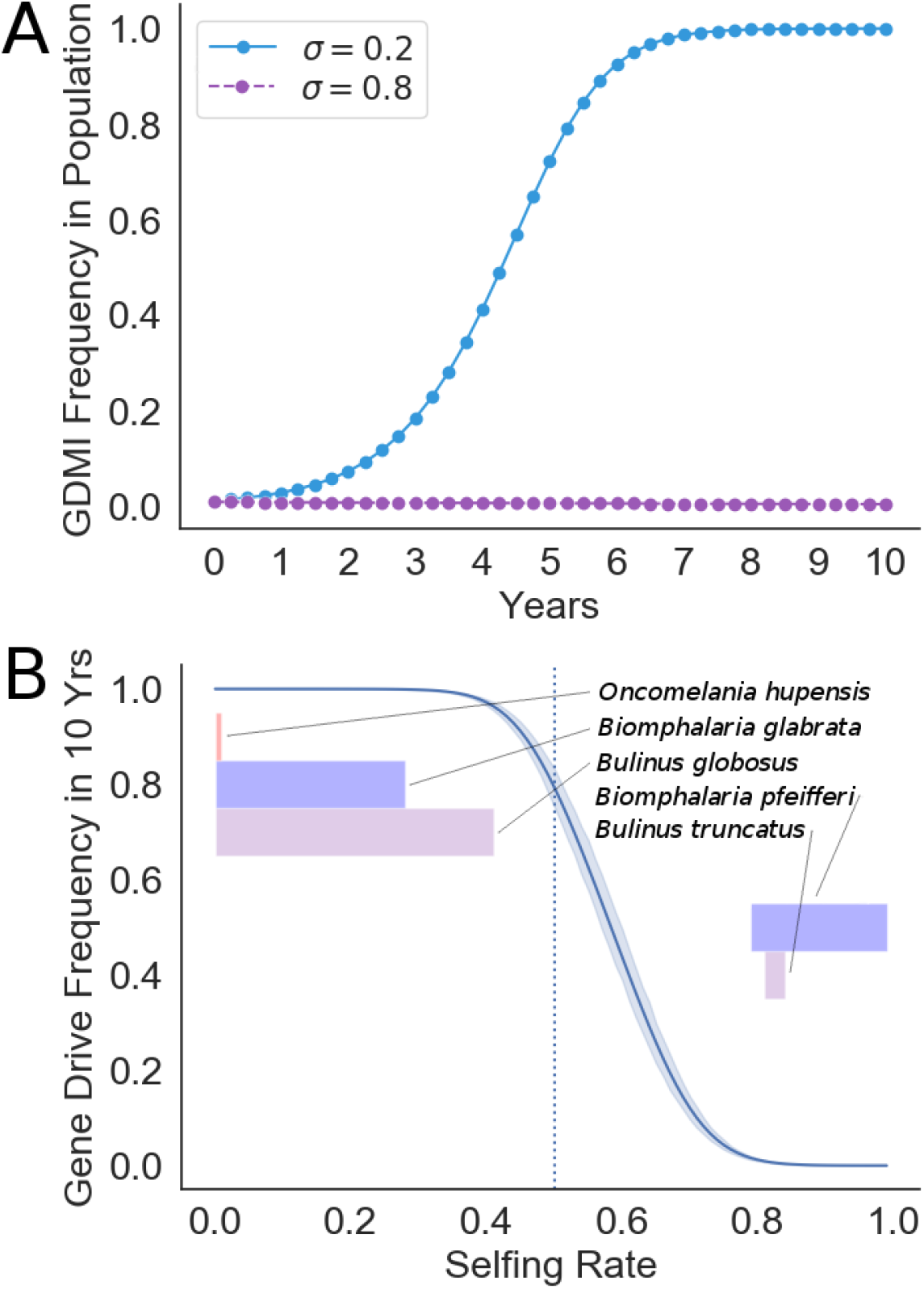
Self-fertilization rate strongly affects establishment of gene drive in a 10 year window. (A) Simulation of gene drive invasion under default conditions when self-fertilization rate is low at 0.2 and high at 0.8. (B) Endpoint sensitivity analysis depicting the gene drive frequency in the population after 10 years under variable self-fertilization rates (σ) from 0 to 1. 95% confidence intervals are reported on the range of results when the fecundity cost of inbreeding varies on the uniform distribution [0,0.6]. The vertical dotted line designates σ = 0.5, which is the value used in future simulations to represent an intermediate selfing rate from those observed. Shaded bars colored by genus display ranges (mean ± 1 s.d.) of observed selfing rates for each host snail species for which empirical measures exist [37]. Vertical positioning of the bars is ordered by minimum selfing rate according to the displayed ranges.

Drive success depends on features in the snail-human-schistosome system beyond selfing. Features intrinsic to the design and deployment of the drive like homing efficiency, fitness cost of the payload, resistance evolution, and the number of releases of GDMI individuals are more easily modified than extrinsic factors which are dependent on the ecological and environmental conditions – size of the snail population, force of infection from the human population, gene flow, and standing genetic variation. Yet the success of GDMI may be sensitive to any of these factors. We explore via endpoint sensitivity analyses how variation in these factors alters the frequency of GDMI after 10 years.

Like results from previous modeling and laboratory studies, we find that homing efficiency has a dramatic impact on the outcome of the gene drive release in a focus population. Under the range of selfing scenarios, low homing efficiency leads to minimal gene drive establishment. Laboratory work in mosquitoes and mice shows homing efficiency above 0.4 is achievable and often exceeds 0.9 [5, 40]. In this range, diminishing returns are observed when H¿0.5 (Fig. 3A). The fitness cost of the genetic payload is not often empirically measured, though for this modeled system, gene drive success is highly sensitive to this parameter: GDMI can establish only when the cost is below 0.4 per gene drive copy in the genome (Fig. 3B). Results improve nearly linearly below a fitness cost of 0.3 per copy. In natural and laboratory populations, resistance to the gene drive mechanism can evolve quickly without the presence of multiple gRNA or selection against resistance formation [41]. Resistance can evolve more quickly when associated fitness costs of the gene drive phenotype are high. The reported mechanisms of resistance are spontaneous mutation and non-homologous end joining which render the Cas9 cleavage site unrecognizable [42]. We combine these associated mechanisms and display the scenarios for the likely range of summed rates of both processes. In a 10-year time frame, a rate of resistance formation greater than 0.2 per meiotic event makes GDMI establishment infeasible (see also SI Fig. S1). With the exception of the deployment strategy, in which the number of releases does not significantly alter establishment drive success, intrinsic factors to the design of the gene drive construct bear heavily on the outcome of GDMI in 10 years.

**Figure 3:**
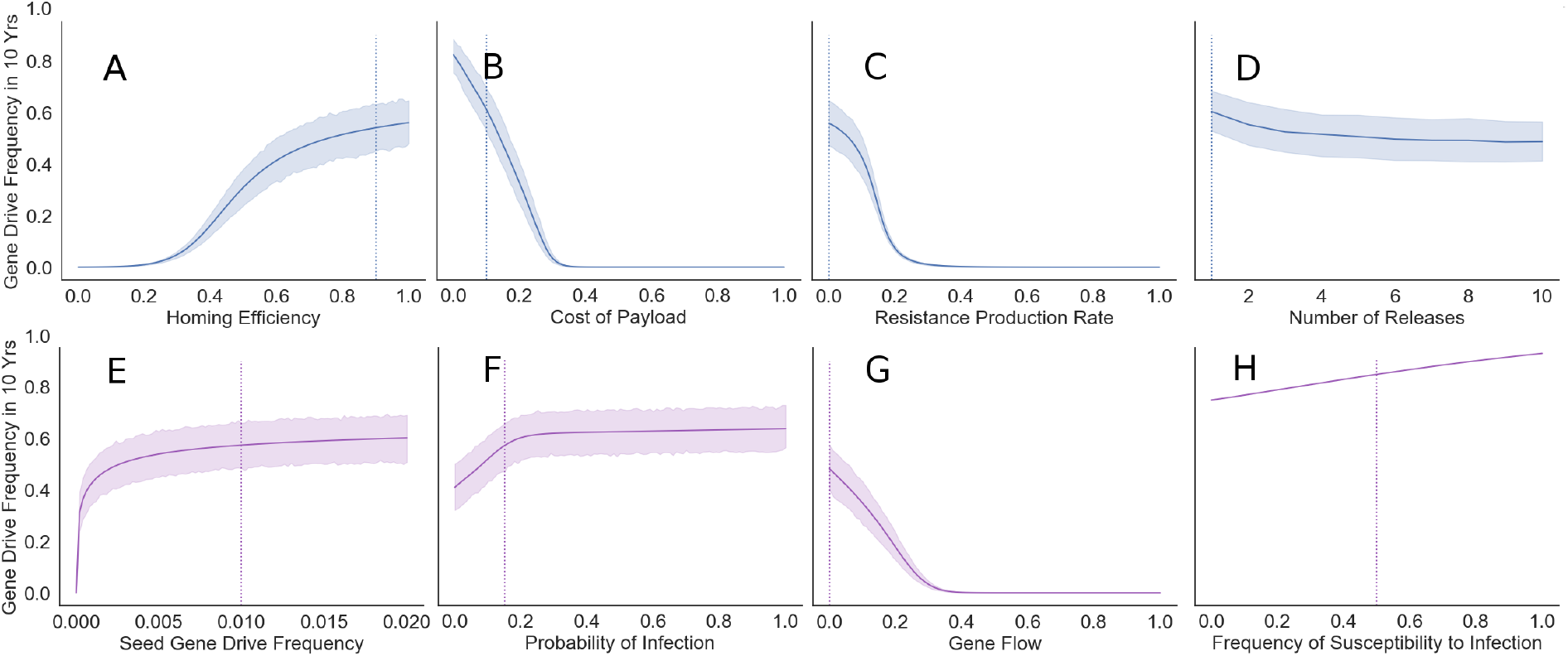
GDMI efficacy across a wide range of endemic conditions and genetic design. Boot-strapped 95% confidence intervals of the mean are reported on the range of results when the self-fertilization rate varies on the uniform distribution [0, 1]. Intrinsic factors to the design and release that may be modified are in the top row in blue: (A) Gene drive homing efficiency (homing success per meiotic event), (B) fecundity cost of the genetic payload (relative fitness loss), (C) rate of production of gene drive resistant mutants (resistant mutants per meiotic event), (D) number of annual GDMI partial releases needed to achieve the size of a single maximum release (e.g. 5 = 1/5th release in each of the first 5 years). Extrinsic factors that are not readily modified are in the bottom row in purple: (E) seed frequency (i.e. proportion introduced in a single release at *t* = 0) of the gene drive engineered snails in the population, (F) the probability a susceptible snail is infected in a generation, (G) bidirectional gene flow as the proportion of the focus population that is replaced by an external source each generation, and (H) starting frequency of snails susceptible to infection in focus the population. Because the frequency of the susceptible genotype is a function of selfing rate, bootstrapping is not appropriate for panel H. Vertical dotted lines depict the default parameter values used in the other figures. Note that the solid line represents a mean across uniformly distributed selfing values, σ [0, 1], rather than the predicted results at σ = 0.5.

We also investigated four extrinsic determinants of GDMI establishment: seed gene drive frequency, force of infection, gene flow, and the standing innate immunity in the snail population. Seed frequency is critical to gene drive spread when only low seed frequencies are possible (Fig 3E). Seeding greater than 1% gives strong diminishing returns. As focus snail populations can vary between hundreds and hundreds of thousands of individuals, this implies that anywhere from one to thousands of snails will need to be raised for a successful introduction, and the size of the focus population will determine the feasibility of release. Similarly, diminishing returns are seen as the probability of infection in a generation increases past 0.2 (Fig 3F). This indicator of endemicity provides the positive selection necessary to propagate the drive in the snail population, as susceptibility to infection is disadvantageous. These results suggest that success is similar for localities experiencing moderate or high burden of disease. Loss of drive alleles from the focus population due to migration inhibits establishment of the drive (Fig 3G). Levels of gene flow greater than 40% (i.e. 40% of focus population alleles are replaced by alleles from a non-evolving background population each generation) bring the drive alleles to undetectable levels assuming immigrants to the focus population lack gene drive immunity. Importantly, GDMI to schistosome infection acts by elevating the level of naturally occurring innate immunity in the snail population. This co-occurring immunity is positively selected under the same conditions as GDMI. Susceptibility is positively selected with weak force of infection due to the fitness costs via reduced egg viability associated with immunity, and immunity is positively selected with moderate to high force of infection due to fitness costs via parasitic castration and reduced lifespan in infected snails [43–45]. High levels of natural immunity will slow the growth of gene drive through direct competition, and therefore, higher susceptibility to infection in a population favors gene drive establishment (Fig 3H). Natural immunity is inherited more slowly, though fecundity for naturally immune snails is assumed higher than GDMI due to added costs of maintaining the genetic payload of the drive.

Although mass drug administration (MDA) is capable of temporary reduction in morbidity, MDA alone is incapable of local elimination at high transmission sites. In these conditions, gene drive offers a potentially promising avenue for coincident MDA and environmental treatment of schistosomiasis. We evaluate the consequences of applying GDMI snails to a community with concurrent annual MDA treatment. We compare the observed reduction in mean worm burden (MWB) between three treatment regimes: gene drive immunity, MDA, and concurrent application of both (Fig. 4). Simulations are conducted under the same default conditions evaluated above with the difference that human-to-snail force of infection is a variable that is determined by the number of mated worm pairs in the human population (SI table 2). The pre-treatment prevalence of infection is assumed to be 80%. The snail-to-human force of infection is a function of the number of infected snails at a water access site, a quantity that diminishes as immunity to infection increases in the snail population.

**Figure 4:**
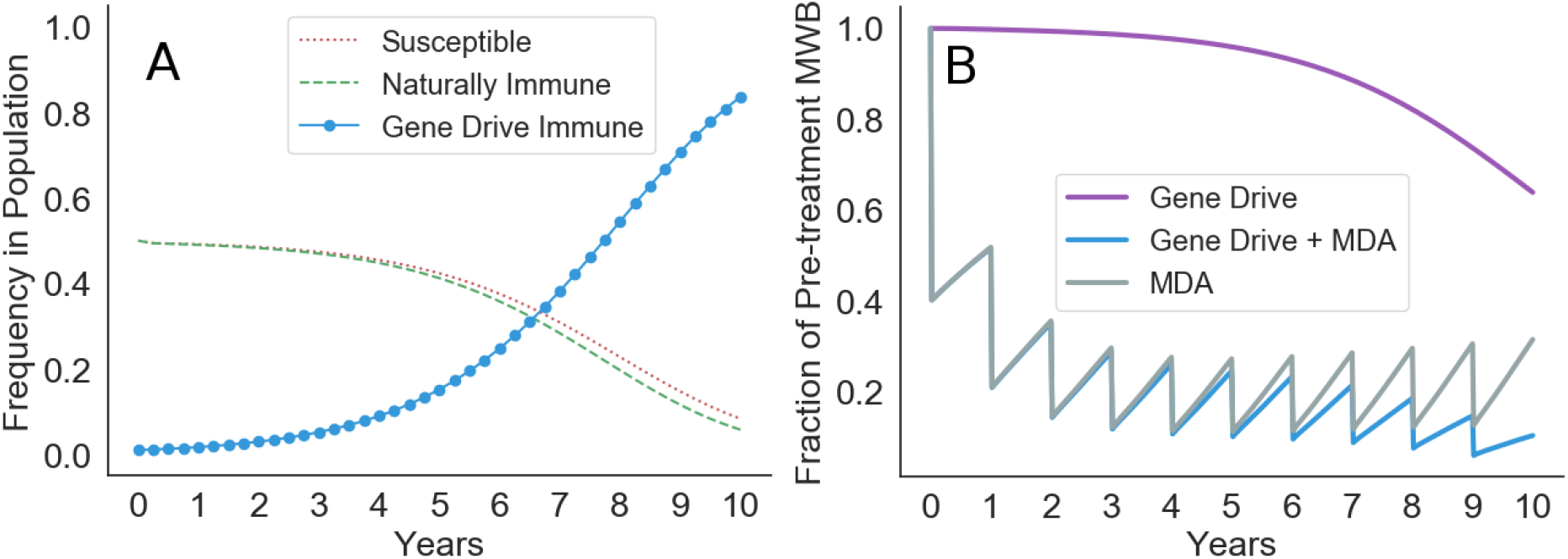
Combining gene drive with mass drug administration. (A) 35% reduction in MWB is observed in 10 years with gene drive alone. (B) Targeted administration of MDA at 60% annual reduction (efficacy*coverage) in MWB results in more rapid but temporary reduction than the use of gene drive. Sustained reductions are achieved with coincident MDA and gene drive treatment.

With gene drive treatment alone a 35% reduction in MWB is observed due to the reduced establishment of new worms in humans and natural mortality of existing adult worms with average lifespan of nearly 5 years [46]. Elimination could be achieved with successful gene drive treatment alone, though the lifespan of adult schistosomes precludes rapid elimination (30 years required for 99% reduction, SI Fig. S14). With annual reduction of MWB of 60% through targeted MDA, alleviation is possible, but elimination is infeasible due to the persistence of infected snails in nearby water access sites. Moreover, immunity in snails wanes due to decreased force of infection on the snail population, resulting in an upward trend in MWB from year 3 onward. Concurrent treatment targeting both snail and human hosts leads to sustained elimination provided resistance formation is low. This is true even when MDA is ended after 10 years because GDMI has reached fixation in the focus snail population (SI Fig. S12-S14).

## Discussion

Our results demonstrate that successful establishment of immunity within a 10 year evaluation period is possible for species of snails with low to moderate selfing rates. Snails species like *B. Pfeifferi* and *B. truncatus*, which are known to self-fertilize at high rates, are likely not desirable targets for GDMI. Many other snail species self-fertilize at lower rates, providing more opportunity for GDMI control of schistosomiasis [25]. Likewise, propensity to self-fertilize can also vary by environmental condition. Panmictic and stable snail populations favor out-crossing, which increases the rate of inheritance of GDMI. This work indicates that the potential for success of GDMI could be evaluated prior to program implementation through genetic studies quantifying selfing rates (e.g. with F-statistics) in intervention areas. In areas with sympatric snail species with differing selfing rates, quantifying the relative abundance of each species and their respective contributions to schistosomiasis burden will inform the potential for success of GDMI locally.

GDMI establishment is sensitive to genetic design and less sensitive to standing genetic variation for immunity. Low payload fitness costs and homing efficiency greater than 50% are essential. Reducing the evolution of resistance to the drive with multiple gRNAs [41] or through other techniques can moderately improve success in a 10 year window and has stronger implications for success after 10 years. Alternative designs incorporating ‘daisy chain’ inheritance or other drive decay mechanisms can provide safeguards to gene drive release in natural ecosystems, and peak GDMI frequency would be contingent on the strength of this decay, which occurs more quickly with fewer loci in the chain [23]. Selfing requires more loci in a daisy chain to achieve high peak frequencies of GDMI prior to decay, therefore this technology also will perform best for preferentially outcrossing species but will be ineffective for large snail populations (SI Fig. S2). Other genetic features like dominance, penetrance, and epistatic interactions are significant considerations for choosing appropriate gene targets (SI Fig. S4-S6). Although optimizing genetic designs is not trivial [47], because modifying snail habitat on a large scale is more challenging, efforts to improve drive construct designs will yield higher returns in successful establishment of GDMI.

Ecological factors that dilute the frequency of gene drive in a focus population (e.g. at a water access site), such as high gene flow due to snail migration or a large snail population size, inhibit timely establishment of gene drive mediated immunity (see also SI Fig. S8). These results indicate that success is most easily achieved in isolated water bodies with smaller snail populations. Snail population sizes in many areas fluctuate dramatically by season [48], therefore introduction of GDMI snails may be best timed when population sizes are at their lowest and have maximum growth potential. Otherwise, GDMI will establish slowly in populations experiencing high seasonality (SI Fig. S10). Populations with shorter generation times will achieve greater GDMI frequencies within 10 years (SI Fig. S7). Future studies should build on this foundational simulation by considering snail migration and water flow between locations to assess whether GDMI snails would be effective in a wider range of scenarios.

There are some caveats and complexities that we have not addressed here. This model was built on the assumption that the gene drive works, i.e. that the gene drive is effective in producing snails that are immune to schistosome infection. While immune alleles associated with the PTC I and II gene clusters in *B. glabrata* can be rapidly selected in experimental conditions, these alleles have not yet been successfully deployed in a gene drive construct. Further genetic work is required to discover gene drive targets in other snail host species. In addition, we have not considered potential interactions with other trematode species, as snails can be the intermediate hosts to species other than schistosomes [49]. Interactions between schistosomes, other trematode species, and immunity can shape the fitness landscape in which GDMI operates, and therefore require further investigation to gauge whether the efficacy of this gene drive approach is sensitive to these interactions. This equally applies to interactions that involve schistosome subtypes that may evade a single GDMI design. Field work to identify sympatric schistosome subtypes will be necessary to evaluate local deployment of GDMI.

These results indicate that the use of GDMI together with MDA could contribute to a longer-lasting reduction of worm burden than either GDMI or MDA alone. This emphasizes that gene drive is one potential tool among several that are currently available, and optimal use would likely be in conjunction with current control methods. GDMI is much more targeted than molluscicides, as it does not destroy the populations of snails and other aquatic life, and thus may be preferred by many stakeholders. Moving forward, it will be necessary to model how gene drives could interact with the variety of other control methods, to assess the optimal combination of methods and timing that would result in sustained elimination.

Modeling is crucial to understand the feasibility of implementing a new technology like gene drive, particularly in a natural system. Although this technology represents a new frontier for controlling disease, pests, and invasive species, the spread of designed genes in a natural setting can carry serious ethical and practical implications [36, 50]. It is therefore prudent to begin any considerations with *in silico* and *in vitro* studies, before proceeding to *in vivo*, with earlier steps informing the next. Further, modeling can ground critical deliberations amongst stakeholders by providing realistic predictions for the effects of a gene drive project and provide a useful ability to rapidly perform new simulations to address questions that stakeholders might have for gene drive developers [51]. This model is an advancement towards a biologically realistic simulation which integrates population genetics, epidemiology, and population dynamics, and can serve as a template for future work in gene drive feasibility analysis.

## Materials and Methods

### Population Genetic Model

We present a model of mixed mating strategy and explore a range of observed selfing rates to understand how reproduction strategy influences success of gene drive technology in a natural population. The gene drive model developed embeds a non-stationary Markov process that accounts for natural inheritance patterns as well as gene drive inheritance and fitness differences among genotypes. In contrast to previous gene drive models, which consider a wildtype allele and gene drive allele, we consider an expanded model with two wildtype alleles: susceptible (*A*) and immune (*B*). A third allele, *B_g_*, represents engineered immunity to infection in the form of a gene drive construct. The set of six genotypes formed by these three alleles is Ω = {*AA, AB, BB, AB_g_, BB_g_, B_g_ B_g_*}. Let *P_i_* be the frequency of each genotype where *i* ∈ Ω. Let ***P_t_*** be the row vector composed of genotype frequencies at time (in generations) *t*. We describe the mixed mating system of genetic inheritance with two transition matrices, ***S*** (self-fertilization) and ***T*** (out-crossing), to describe the transitional probabilities from generation *t* to *t* + 1.

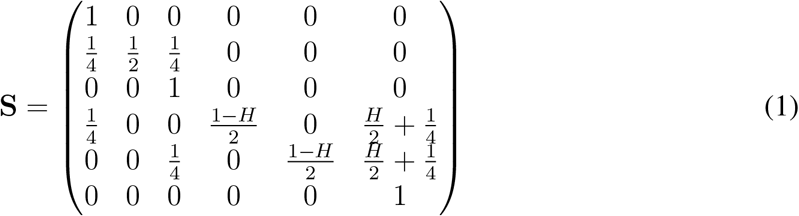

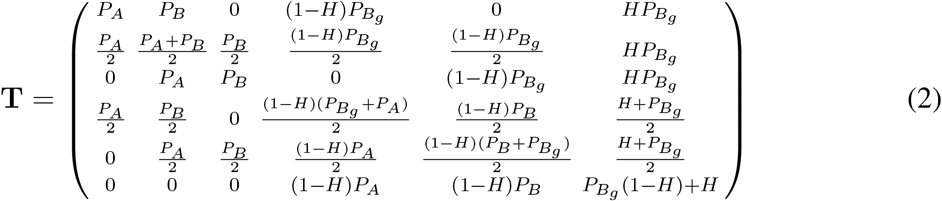

Homing efficiency *H* takes values between 0 (Mendelian inheritance) and 1 (complete fidelity of gene drive mechanism). A matrix ***Q*** can be formed to represent the mixed mating system with self-fertilization rate and cost of inbreeding given by σ, ξ ∈ [0, 1], respectively.

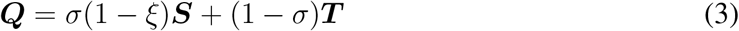

In the absence of population dynamics and fitness differences between genotypes, equation (4) would suffice to describe genotype frequency changes over time.

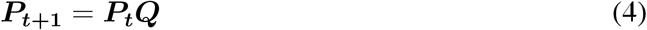

To accurately represent the evolution of traits in the snail-schistosome system, we relax these simplifying assumptions by incorporating fitness differences in response to viability and fecundity selection. We also introduce overlapping generations with density dependent recruitment in the snail population (SI Equns. S1-S45). Model simulations recapitulate laboratory results under the same conditions (SI Fig. S3). We derive analytical solutions at long term equilibrium (SI Equns. S46-S55) and analyze the evolution of resistance to the gene drive mechanism (SI Equns. S56-S58). The model is expanded to simulate the effects of ‘daisy chain’ drive (SI Fig. S2, Equns. S59-S68), and invasion analysis is performed for key variables, while a stochastic model is used to observe extinction conditions (SI Fig. S9, Equns. S69-S71).

### Epidemiological Model

The genetic model is integrated with an epidemiological model of schistosomiasis through the fraction of immunity in the snail population, *ρ*.

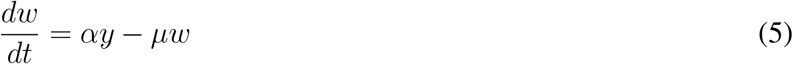

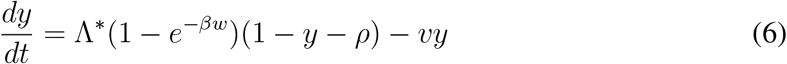

*α* and *β* are transmission rates governing the conversion of snail infection prevalence, *y*, to adult worms, *w*, in humans and vice versa. Gurarie et al. outlined the transmission from humans to snails is saturating with increasing worm burden [52]. The asymptote is the pre-treatment force of infection, Λ*^*^*. *µ* and *v* are the death rates of adult worms and infected snails, respectively.

Equations (5) and (6) are integrated in 3 month intervals, corresponding to the expected generation time of the snail population. The probability of a new infection per susceptible snail in a generation determines the strength of selection for immunity in the genetic model.

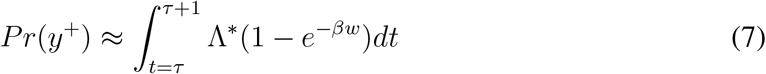

Parameter values and initial endemic conditions are detailed in SI tables 1 and 2. Calculations for equilibrium values and reproduction numbers are made in SI Equns. S72-S91. The epidemiological model was not used to simulate figures 2 and 3. Instead, the probability of new infections was held constant at the equilibrium value calculated for endemic conditions with the integrated genetic and epidemiological model (SI Fig. S11). Figure 4 incorporates both dynamic models to evaluate single and paired treatment. MDA is modeled as instantaneous annual treatment. Percent reduction in worm burden during each treatment was 60% and is the product of coverage and efficacy.

Python code for simulations is available at www.github.com/grewelle/ModelGeneDriveSchisto.

## Acknowledgments

We thank members of the De Leo lab and ECHO laboratory (Ecole Polytechnique Fédérale de Lausanne) for continued support in this work. We also thank John Pringle for insightful comments on this work. REG is funded by the Stanford Graduate Fellowship, ARCS Fellowship, and the Stanford-EPFL exchange fellowship. **Conflicts of Interest**: JT and EKON were seed funded by the Merck Innovation Cup 2016 for research on schistosomiasis, and previously employed as external consultants to the Global Health Institute of Merck (KGaA) which produces treatments for schistosomiasis. REG and GADL were partially supported by the National Science Foundation’s grants DEB-2011179 and ICER-2024383.

## Author Contributions

R.E.G, J.P.S.,J.T., E.K.O.N. conceived the research; R.E.G. performed the research; R.E.G wrote the paper; and R.E.G., J.P.S, J.T. E.K.O.N., C.G.R., G.A.D.L edited the paper.

## Supplement

### Population genetic model

The evolution of a focal population seeded with gene drive mediated immune (GDMI) snails is described in rudimentary form in the main text with transition matrix ***Q***. This describes a system of inheritance where a susceptible and immune allele are present in the population, and the GDMI allele exhibits the same immunity as the naturally occurring immunity. The three alleles form six distinct genotypes in a diploid species. ***Q*** represents random mating with no selection and full population replacement each generation. In a natural system assortative mating, selection, and iteroparity are known to occur. Because assortative mating as a function of innate immunity to schistosome infection has not yet been demonstrated in host snail species, we maintain this assumption in our population genetic model. However, because several modes of selection are described for this immunity and iteroparity produces overlapping generations, other assumptions for ***Q*** must be relaxed to accurately reflect the evolutionary dynamics of the snails. Viability and fecundity selection are separately accounted in their contribution the fitness of each genotype, as their relative importance in determining the rate of evolution changes with the model of population dynamics and replacement.

### Birth-death process

Snail recruitment is density-dependent. Adult lifespan extends past the mean generation time, allowing for nearly continuous reproduction after sexual maturity. We assume that background mortality is density-independent and that the population replacement rate is modulated by changes in mortality rate provided density-dependent recruitment is sufficient to for full replacement. In contrast to a population model described by reproduction proceeded by culling to a carrying capacity, we model snail population dynamics with culling proceeded by reproduction to a carrying capacity. The sub-population size of genotype *i* is described through time with the two step process: (1) death and migration yield the reproducing population of genotype *i*, *N̄*_*i*_(*t*), which (2) give birth to offspring according to equation 2.

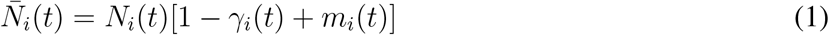

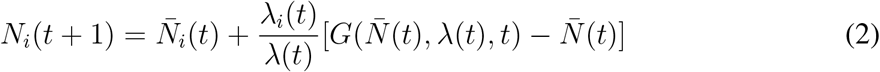

where *N_i_*(*t*) is the genotype sub-population size in generation *t*, *γ_i_*(*t*) is the fractional mortality in generation *t*, *m_i_*(*t*) is the fractional net migration in or out of the focal population in generation *t*, *λ_i_*(*t*) is the partial finite growth of genotype *i* after mortality and migration, and *λ*(*t*) is the maximum total finite growth of the population. Because deaths are separately accounted in this birth-death process, here *λ*(*t*) resembles fecundity (i.e. when mortality is absent, fecundity and finite population growth are equivalent). *G*(*N̄*(*t*)*, λ*(*t*)*, t*) is the discrete time population growth function which describes total population growth. We use a logistic growth function in the simulations throughout the main text and supplement.

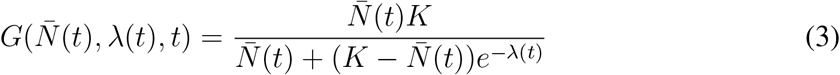

### Viability selection

Viability selection on immunity to schistosome infection is incorporated in the fractional mortality term, *γ_i_*(*t*), which is a function of background mortality (neutral) and loss from the reproductive population through infection (directional selection), which has been demonstrated to substantially increase mortality and castrate snails. For these two reasons, we assume that infected snails are removed from the reproducing population. Reproductive compensation has been observed for snails exposed to miracidia, whereby these snails produce offspring at higher rates following exposure. This could counter lost reproduction after infection. However, a genetic link between reproductive compensation and immunity has not yet been shown. Therefore, we assume that the mechanism of reproductive compensation is phenomenological, being linked to exposure but not patent infection, rather than linked to host genetics, and does not produce fitness differences between genotypes. Explicitly, we write *γ_i_*(*t*) as

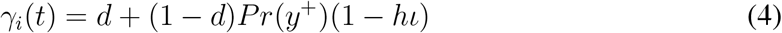

*d* is the adult background mortality per snail in a generation. *P r*(*y*^+^) is the per snail probability of acquiring a new infection. *h* is the dominance coefficient, which takes values between 0 and 1, with a value of 0.5 indicating co-dominance and a value of 1 indicating complete dominance of the immune allele over the susceptible allele. ι ∈ [0, 1] is the degree of immunity conferred by the immune allele compared to the susceptible allele. Genetically, this value can be equated to the penetrance of innate immunity. A value of 0 indicates no additional immunity, while a value of 1 indicates full immunity. The loss of reproductive individuals from the population contributed by infection compared to the background mortality determines the strength of viability selection for immunity in the population.

### Fecundity selection

Inbreeding and immunity are known to negatively impact reproductive success in host snail populations. Inbreeding can reduce fecundity and egg viability, while immunity is associated with low egg viability [53, 54]. Because the population model tracks reproductive individuals, and recruitment of offspring to the reproductive class is density-dependent, offspring viability can be treated as a component of fecundity. Using the broad definition of fecundity as the offspring surviving to adulthood, we institute fecundity costs for inbreeding by self-fertilization and for maintenance of immune alleles. Cost of immune maintenance, *C*, is directly related to the phenotype and is dose-independent (i.e. 2 immune alleles are not more costly than 1 immune allele with full dominance). An additional cost, *C_g_*, is associated with maintenance of the genetic payload in the drive construct. This cost is dose-dependent; gene drive homozygotes carry a two-fold cost compared to heterozygotes. Let the set of alleles {*A, B, B_g_*} be indexed as {1, 2, 3}. The fecundity of each genotype, *f_i_*, can be represented as:

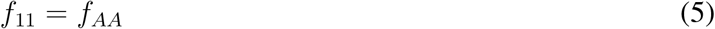

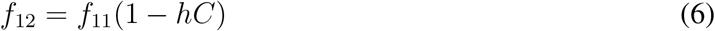

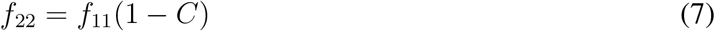

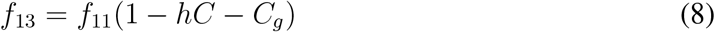

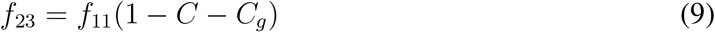

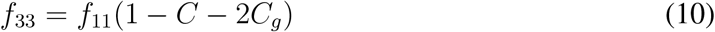

The inbreeding cost is not directly associated with a specific genotype, but rather is incorporated as ξ ∈ [0, 1] in the calculation of ***Q*** (equation 3 main text):

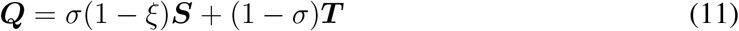

Although *ξ* is a cost not directly applied to any genotype, because it is a cost of inbreeding due to self-fertilization and self-fertilization produces homozygotes with higher frequency than outcrossing, this cost reduces the fitness of homozygotes relative to heterozygotes when selfing is common in the population. Inbreeding is assumed to only affect the F1 generation of selfing parents, and associated costs are not separately tracked through descent in future generations. This treatment is reasonable for a sufficient degree of outcrossing which mixes lineages in the population. Highly inbred snail populations are shown to be insensitive to inbreeding depression caused by selfing, presumably due to purging of deleterious alleles from the gene pool, and therefore inbreeding costs are low in the absence of outcrossing. Table S1 gives parameter values used in the genetic model, many of which are known to vary by species or even by population and environmental conditions. Values were chosen to be centered in the range of observed values with references given to empirical measurements. Results can differ by system, and the endpoint sensitivity analyses in Fig. 2 and 3 of the main text provide indication of the most sensitive parameters.

**Table 1:** Default parameter values for the genetic model

| Parameter | Description | Value | Ref. |
| --- | --- | --- | --- |
| $d$ | background mortality per generation | 0.5 | calculated with reference to *B. pfeifferi* [55, 56] |
| $h$ | dominance coefficient for immune allele | 1 | supported by Fig. S3 and [57] |
| $Pr(y^+)$ | probability of infection per generation | 0.15 | derived from epidemiological model (equ. S83) |
| $\iota$ | penetrance of immunity | 0.8 | [53] |
| $C$ | cost of immunity | 0.2835 | model-derived to yield symmetric selection for and against immunity, stabilizing the phenotypic ratio at 1:1 |
| $C_g$ | cost of payload per copy | 0.1 | [58] variable/theoretical |
| $\xi$ | cost of inbreeding | 0.3 | [59] highly variable |
| $H$ | homing efficiency | 0.9 | [60] |
| $\sigma$ | selfing frequency | 0.5 | [59] highly variable |
| $f_{AA}$ | per generation fecundity of a susceptible snail | 20 | [53] |
| $P_{AA}(t=0)$ | natural initial frequency of susceptible genotype | 0.5 | [53, 57] |
| $P_A(t=0)$ | natural initial frequency of allele A | $\frac{\sigma - \sqrt{16P_{AA} - 24\sigma P_{AA} + \sigma^2(1+8P_{AA})}}{4(\sigma-1)}$ | calculated for a given $\sigma$ |
| $P_B(t=0)$ | natural initial frequency of allele B | $1-P_A$ | calculated for a given $\sigma$ |
| $P_{AB}(t=0)$ | natural initial frequency of heterozygote | $\frac{4P_AP_B(1-\sigma)}{2-\sigma}$ | calculated for a given $\sigma$ |
| $P_{BB}(t=0)$ | natural initial frequency of immune genotype | $\frac{P_B^2 + P_AP_B\sigma}{2-\sigma}$ | calculated for a given $\sigma$ |

The Markov process described in the main text with equations 3 and 4 represents a semelparous population that reproduces once per generation, and adults are completely replaced by offspring with no consideration for fitness differences between genotypes. Instead, however, snail host populations reproduce continuously with overlapping generations. We devise a modified Markov process to describe these evolutionary dynamics, incorporating the fitness differences detailed above in equations S5-S10. We consider the modified transition matrices:

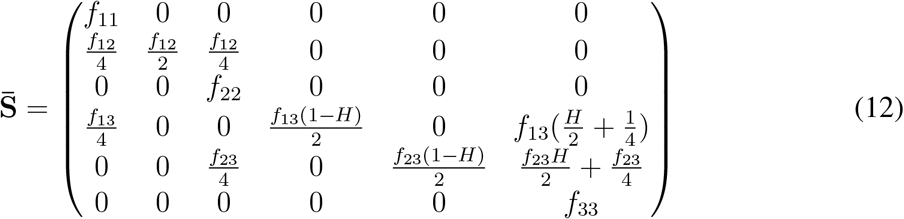

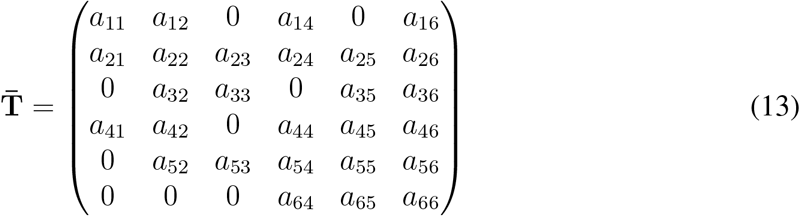

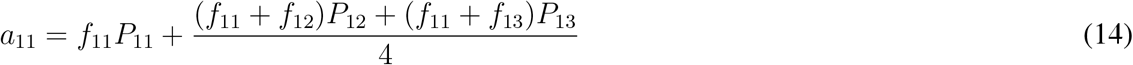

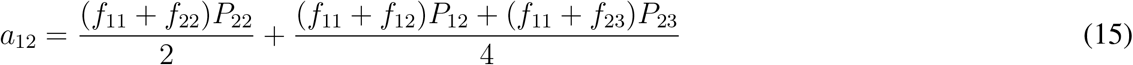

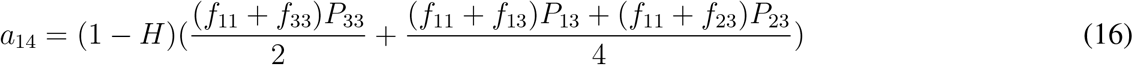

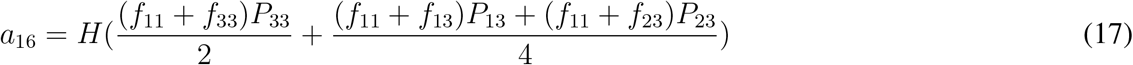

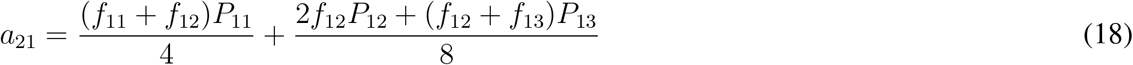

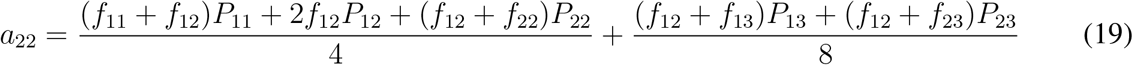

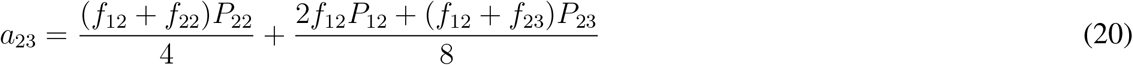

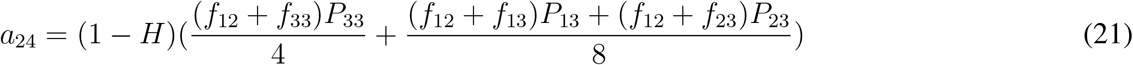

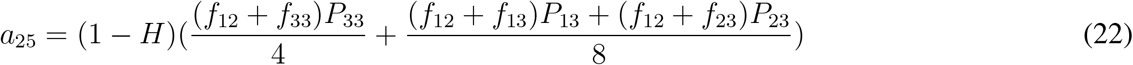

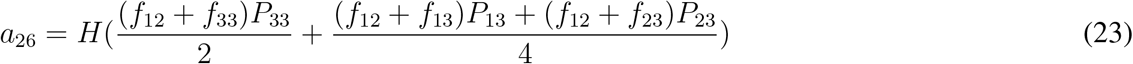

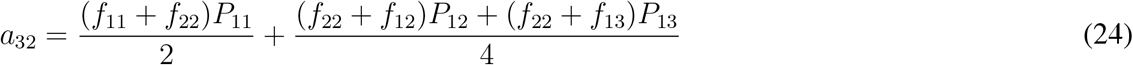

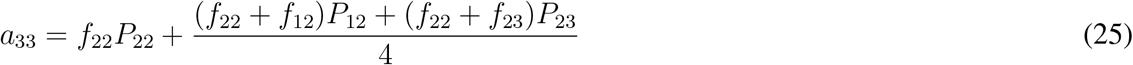

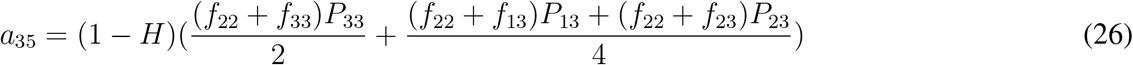

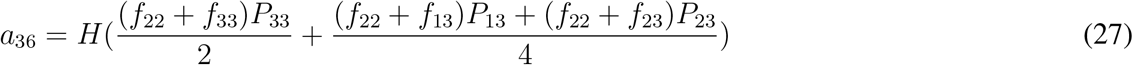

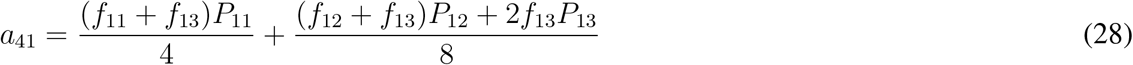

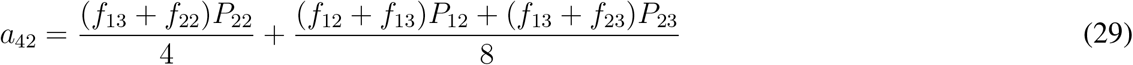

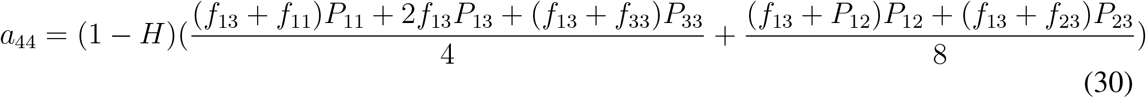

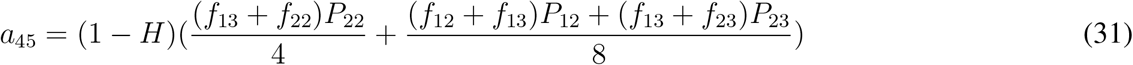

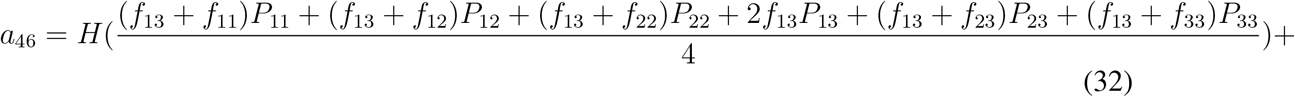

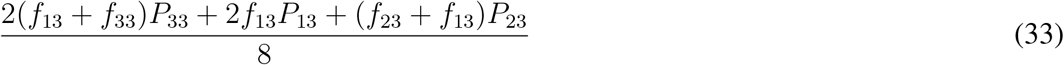

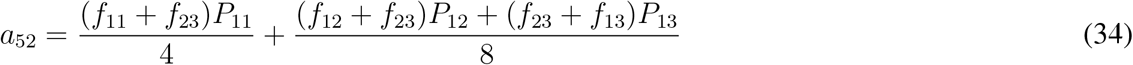

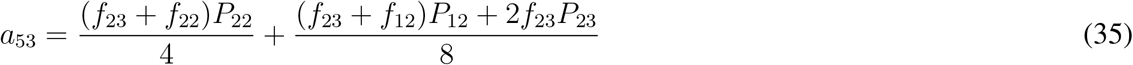

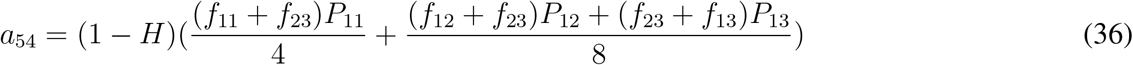

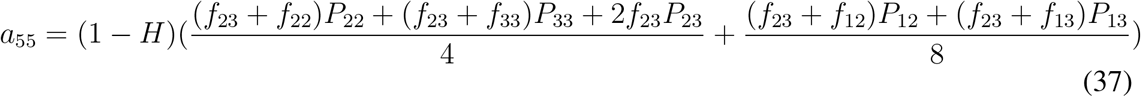

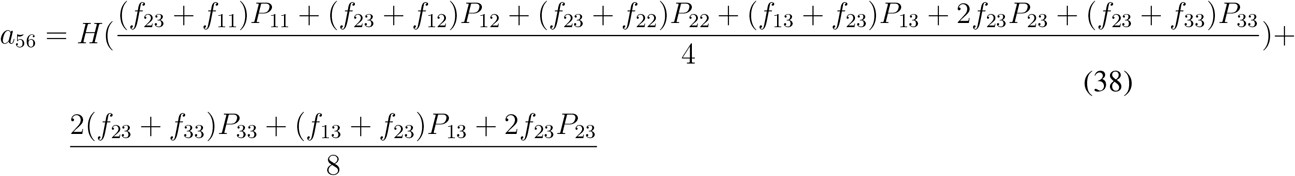

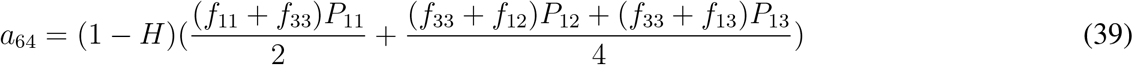

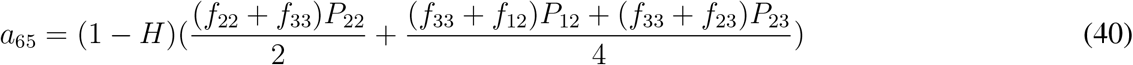

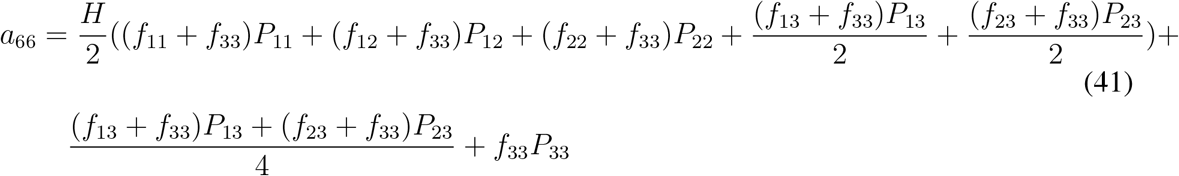

Modifying the equation for ***Q*** in the main text to reflect the incorporation of demography and fitness differences between genotypes, we achieve:

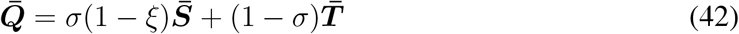

The vector of genotype frequencies at time *t* is denoted ***P*** (*t*). *P_i_*(*t*) represents the frequency of genotype i at time *t*. The vector of partial growth rates is ***λ***(*t*):

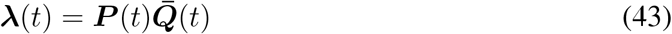

and the sum of the elements of this vector is *λ*(*t*):

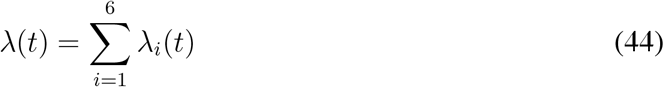

These values can be substituted in equation S2 to track the evolution of the population.

### Establishing initial genetic conditions

Prior to deploying GDMI snails in a naive population, the standing background genetic variation for susceptibility to infection has some influence over the success of GDMI. The two forms of genetic variation that are important to consider are: the frequency of susceptibility and the distribution of the susceptible allele across the genotypes. High self-fertilization frequencies favor homozygous populations, which exposes the susceptible allele to selection (assuming it is recessive). Several studies have measured susceptibility empirically through challenge experiments in laboratory conditions. Snail populations that are not far removed from a natural parental lineage demonstrate intermediate levels of susceptibility, though these results vary with miracidial dosing. For simplicity we fix the standing natural frequency of susceptibility at 0.5. The immune phenotype is, therefore, at 0.5 frequency as well in our idealized starting conditions. In a mixed-mating system, which these snails exhibit, the distribution of the *A* and *B* alleles across the three genotypes is modulated by selfing frequency. The transition matrix describing the evolution of the three naturally occurring genotypes is

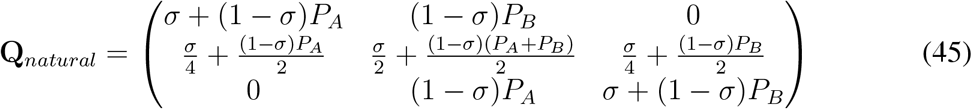

Here we denote *P_A_* and *P_B_* as the allele frequencies, and *P*_11_(*t*), *P*_12_(*t*), *P*_22_(*t*) as the genotype frequencies at time *t*. To establish initial genetic conditions, we find the equilibrium genotype frequencies given a frequency of self-fertilization. We assume that the genotype frequencies are the result of selection but that otherwise selection is not stronger than the equilibrium behavior of the transition matrix under neutral conditions. Therefore, genotype frequencies can be solved given the frequency of the susceptible genotype. Equilibrium behavior of this transition matrix is strong, with genotype frequencies approaching equilibrium geometrically so that equilibrium is effectively reached within 2 generations. In the absence of imposed selection, the allele frequencies remain constant through each generation despite the changing genotype frequencies. This is the reason genotype frequencies are represented as functions of time above, while allele frequencies are not. From matrix multiplication, we know the frequency of the susceptible genotype in the next generation using **Q***_natural_* is:

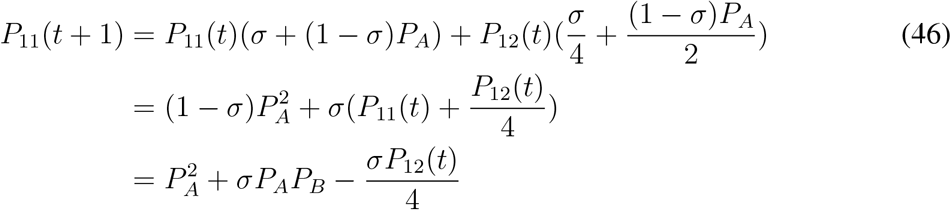

Solving the difference equation yields:

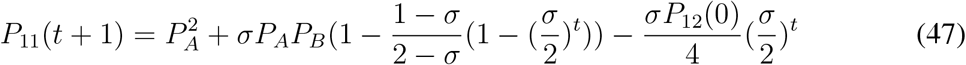

The limiting distribution of genotype frequencies can be solved as *t → ∞*. For the susceptible genotype, this gives:

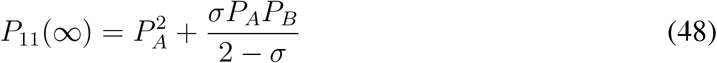

The same derivations can be performed for the other two genotype frequencies from the transition matrix to achieve the limiting distribution.

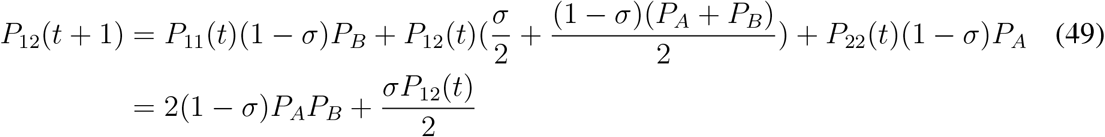

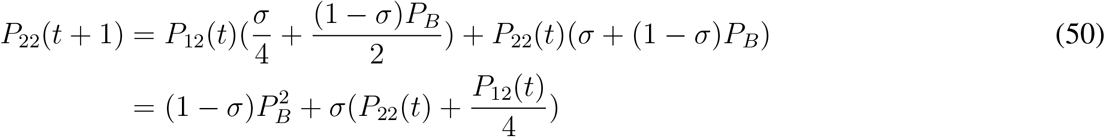

The equilibrium values for the heterozygote and immune homozygote are:

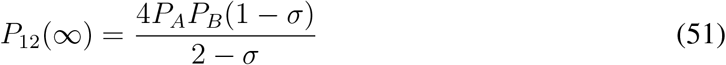

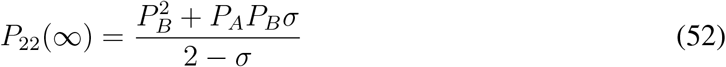

These results differ from the results presented by Karlin [61] due to a presumed typographical error in his text. A population with 50% susceptible genotype at equilibrium is assumed when GDMI snails are introduced (simulation *t* = 0). Given that *P*_11_(*t* = 0) = 0.5, *P_A_* can be solved:

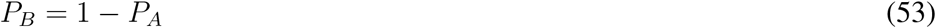

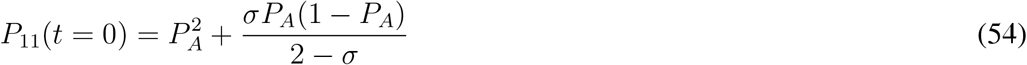

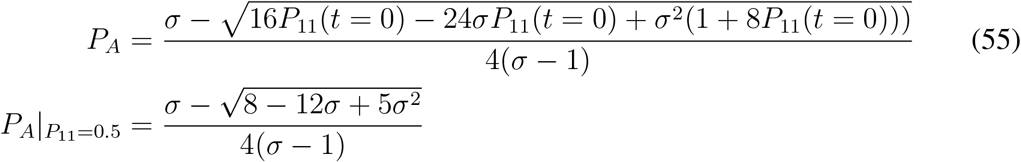

### Evolution of resistance

Resistance to the drive mechanism can readily develop if the target sequence on the homologous chromosome is mutated so as to be unrecognizable by guide RNA. This occurs primarily through non-homologous end joining (NHEJ), which is an alternative mechanism of double strand break repair that can occur instead of homology directed repair. Point mutations and standing genetic variation at the target locus can also result in resistance to the drive mechanism. Because NHEJ is a common repair pathway in most organisms, it is the primary producer of resistance in the population, especially as the drive construct increases in frequency in the population. Resistance formation via this pathway occurs due to misrepair after cleavage from the Cas nuclease and occurs proportionally to the number of cleavage events that occur in the population each generation. Homing efficiency is a function of the predominance of homology directed repair over NHEJ, though not every failed drive event is due to NHEJ. Homing efficiency at the population level declines as resistant alleles accumulate. Resistant alleles may represent a spectrum of mutations, and separately accounting for the variety of alleles and their respective fitness requires exponential expansion of the number of genotypes tracked in this genetic model. To simplify, we assume that resistant alleles are equivalent in fitness to their natural counterparts. This gives them a fecundity advantage to the drive allele, which carries an additional cost due to the genetic payload. In a randomly mating population, outcrossing events will randomly pair resistant alleles with each other, natural alleles, or the drive allele. The consequence of non-assortative mating is that the formation of resistant alleles is proportional to the number of gene drive heterozygotes produced due to failed homing. The number of gene drive heterozygotes (hybrids) produced each generation is:

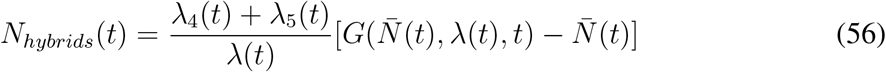

The homing efficiency in generation *t* can be calculated from the maximum homing efficiency without NHEJ, *H*_0_, at *t* = 0 when resistant allele accumulation is lowest and determined only by background resistance due to standing genetic variation.

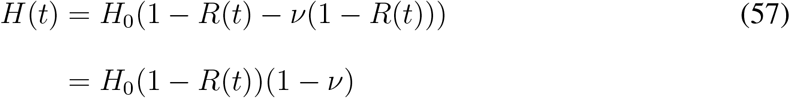

*R*(*t*) is the frequency of resistant alleles in the pool of natural and resistant alleles (excluding the drive allele). *ν* is the per homing event rate of production of resistant alleles. If the population is in mutation-selection-drift balance, the rate of production is almost entirely due to NHEJ. The rate of accumulation of resistant alleles is the fraction of the total gene drive heterozygotes (hybrids) produced that are resistant multiplied by the fraction of hybrids produced in the total population each generation.

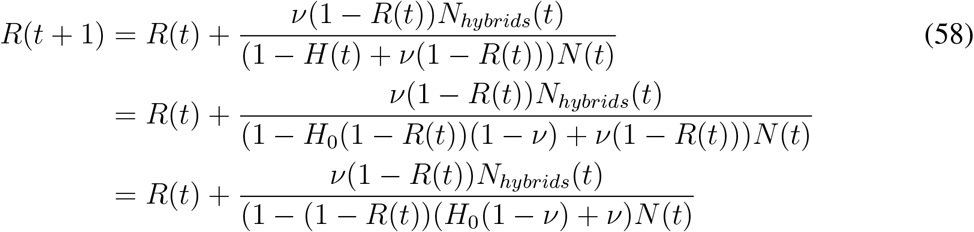

Should mutations leading to resistance be deleterious compared to the respective naturally occurring alleles, resistance is expected to spread more slowly in the population than is depicted in Fig. 3 of the main text. Conversely, should mutations be beneficial, spread will occur more rapidly. The cost of resistance can be modulated by choosing neutral regions for lower cost and regions under strong selection for higher cost (e.g. ribosomal RNA genes). Fitness costs due to off-target CRISPR-induced mutations are possible, but experimental evidence indicates that the frequency of these mutations is low (¡2%). Moreover, these mutations do not exhibit drive and are not related to the evolution of resistance. Off-target mutations can be reduced with properly specific gRNA design and restricted expression of the CAS9 protein. The influence of resistance evolution on the establishment of GDMI is explored in Fig. S1.

### Daisy chain loci

Previous authors have proposed mechanisms to safeguard the spread and persistence of gene drives in wild populations. One prominent mechanism is known as a daisy drive, which borrows its name from the concept of a daisy chain in which elements are connected in a series. Daisy drives are split drives which split the drive element from the payload by positioning them on separate loci, ideally on different chromosomes for independent inheritance. Daisy drives incorporate multiple splits, with one drive element necessary to produce super-Mendelian inheritance of the next element in the chain. The base of the chain is a non-drive element and the tip of the chain is the genetic payload containing the gene of interest to be inherited in the population. Each element in the chain increases in frequency temporarily and then decays as the preceding elements decline in frequency in the population through natural selection. Each element in the daisy chain is considered haplosufficient, so one copy produces the intended homing efficiency of the proceeding element. Because daisy chain elements (loci) are introduced together in engineered individuals, spread of the payload gene occurs locally within a lineage in association with other loci and cannot be modeled using the same framework so far introduced which describes random pairing of alleles from one locus. The homing efficiency associated with the payload locus is primarily determined by this local lineage-based process of inheritance of daisy chain loci, especially when loci are at low frequency, and secondarily through outcrossing events with other lineages in the population that maintain daisy chain loci. The secondary process becomes consequential at high frequencies of daisy chain loci in the population. Homing efficiency of the payload corresponds is determined by the frequency of the preceding locus in the daisy chain. We set *δ*(*t*) as the daisy chain coefficient modulating the homing efficiency. The daisy coefficient is directly proportional to the co-occurrence of the payload and the preceding drive element and can be calculated as follows through time for lineage-specific inheritance (i.e. each outcrossing event occurs with a snail with no drive alleles):

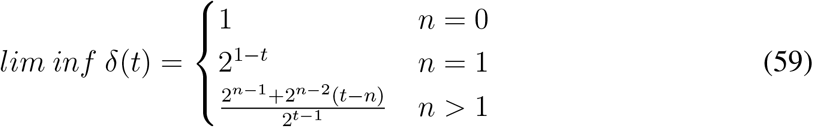

**Figure 1:**
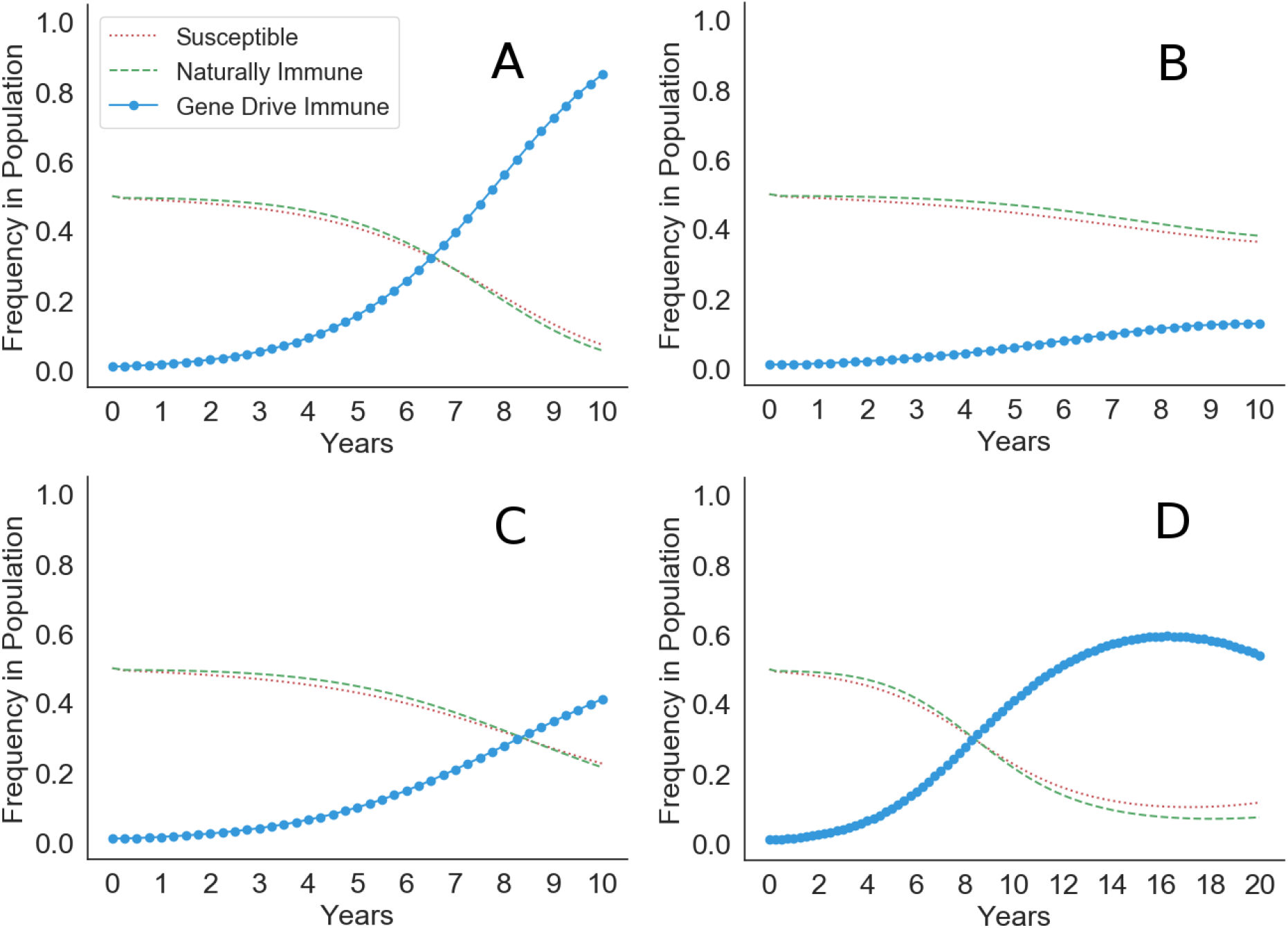
Forward simulations under fixed epidemiological conditions of the spread of GDMI with various resistance production rates per homing event. (A) No resistant alleles are produced. (B) Resistant alleles are produced with 20% of homing events. GDMI achieves only low frequency in the population due to rapid evolution of resistance to the drive mechanism. (C) Resistant alleles are produced with 10% of homing events. GDMI rises slowly, achieving half the frequency in the population compared to conditions where resistance does not evolve. (D) Resistant alleles are produced with 10% of homing events as in panel C. In 20 years it is evident that the frequency of resistant alleles outpaces the homing efficiency benefits in inheritance of GDMI, and GDMI declines after reaching intermediate frequency (eventually to negligible frequency).

*n* is the number of splits in the drive design, which is equivalent to the number of drive elements in the daisy chain (excluding the payload). This is the lower limit for the value of *δ*(*t*), as homing efficiency for the payload increases with the accumulation of the preceding drive element in the population so that with each outcrossing event, the mate outside the primary lineage may carry the preceding drive element. The frequency of the preceding drive element in the population is always lower than the frequency of the payload, and is the frequency of the payload in the prior generation before the peak frequency of the payload is reached. Fitness costs of each drive element are assumed negligible compared to the cost of the genetic payload at the tip of the daisy chain. After the peak frequency is reached, the frequency of the preceding drive element is lower than the frequency of the payload in the prior generation. We can state the following:

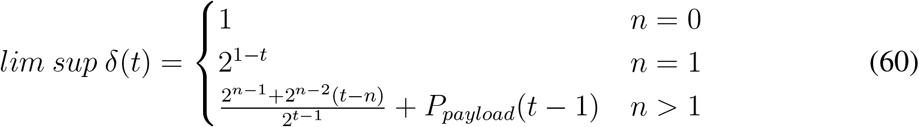

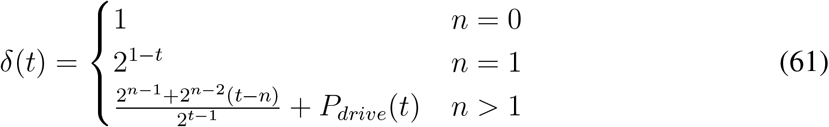

where *P_payload_*(*t*) and *P_drive_*(*t*) are the frequencies of the genetic payload and the associated preceding drive element in the daisy chain, respectively. *lim sup δ*(*t*) = *δ*(*t*) prior to peak payload frequency. Equation S60 is useful to calculate the peak frequency of the payload in a daisy drive system without the need for a system of equations for each locus, which quickly becomes complex with additional elements in the chain. If we consider *H*_0_ as the maximum homing efficiency without a daisy chain design (1 locus, *n* = 0), then the observed homing efficiency through time can be calculated as:

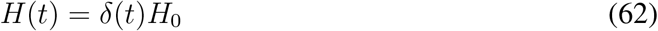

The homing efficiency at time *t* above represents the calculation for non-overlapping generations where the population is fully replaced with offspring each generation. When generations overlap, this homing efficiency underestimates the observed homing efficiency because younger adults reproduce alongside older adults, which maintain higher loads of daisy chain loci from fewer outcrossing events. Older adults produce more GDMI offspring as a result. The observed homing efficiency at the population level at time *t* is a function of the survival of each age class. We calculate the surviving fraction of the payload allele as:

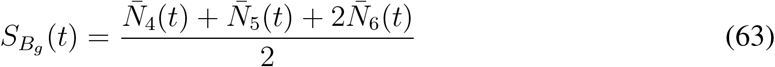

Let ***S_Bg_*** be the vector of surviving fractions of the payload allele through time.

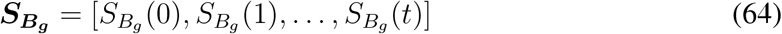

An age distribution, ***ZN̄*_*Bg*_**, can be produced by calculating the survival of each age class through time and normalizing the vector by the sum of the elements.

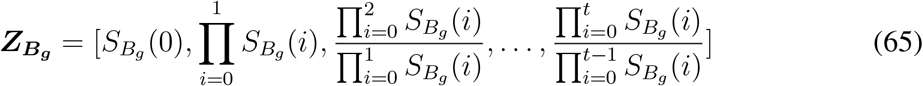

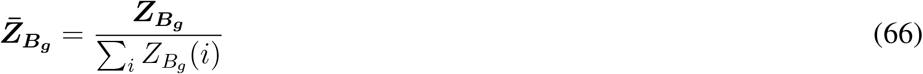

An adjusted daisy chain coefficient, *δ_adj_* (*t*), can be produced by calculating the dot product of the vector of daisy chain coefficients and the age distribution:

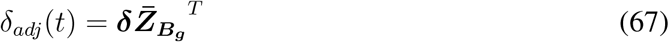

Homing efficiency at time *t* for a population with overlapping generations is therefore:

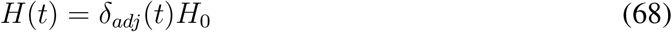

This value for the homing efficiency in each generation can be substituted into the existing framework described to calculate the frequency of GDMI through time. Fig. S2 shows the trajectory of GDMI with the use of an increasing number of daisy chain loci.

### Model validation with empirical data

Tennessen et al. 2015 [57] performed selection experiments using two infection conditions: 10 and 30 miracidia per snail. These snails were 10 generations from natural *Biomphalaria glabrata* breeding populations and were kept together to breed during each generation of selection. After challenging each group of snails with miracidia, infected snails were removed from the breeding population. Selection for immunity was evident and genetically based as given by experimental evidence of decline in infection through the 6 generations of challenges. We modify our model to replicate these experimental conditions, first by assuming that removal from the breeding pool only occurs via infection (mortality = 0). We assumed a high probability of infection of 80% for susceptible snails in the 30 miracidia experimental condition. Because the snails are kept in close proximity, and *B. glabrata* are known to outcross frequently, we assume that outcrossing was the exclusive mode of reproduction (i.e. selfing = 0). Initial allele frequencies were calculated on the basis of the frequency of observed infections (approx. 57%) in the 30 miracidia experimental condition for a probability of infection of 80 % for susceptible snails. GDMI was absent and set to a frequency of 0. Otherwise, parameters were unaltered from simulation conditions in the main text. Initial allele and genotype frequencies were assumed the same between the two experimental treatments, and the probability of infection of susceptible snails was calculated given the frequency of observed infections in the first challenge (48%). The probability of infection for the 10 miracidia treatment is 70%. The curves produced by the model in Fig. S3 of expected infection frequencies given these two calculated probabilities (70% and 80%) reflect the observed data well despite some assumptions (e.g. no self-fertilization) and experimental variability. We consider this fit qualitatively similar because some unknown experimental conditions are assumed, and therefore represent one of the possible model outcomes. However, empirical evidence suggesting that immunity is a dominant trait and that it is regulated by a gene complex, which is tightly linked, corroborates our use of a one locus, complete dominance (*h* = 1) model. Additionally, model parameters used in the simulations of the main text are able to generate similar evolutionary dynamics to experimentally achieved evolution, and therefore, their values are further supported by our results in addition to support from literature.

**Figure 2:**
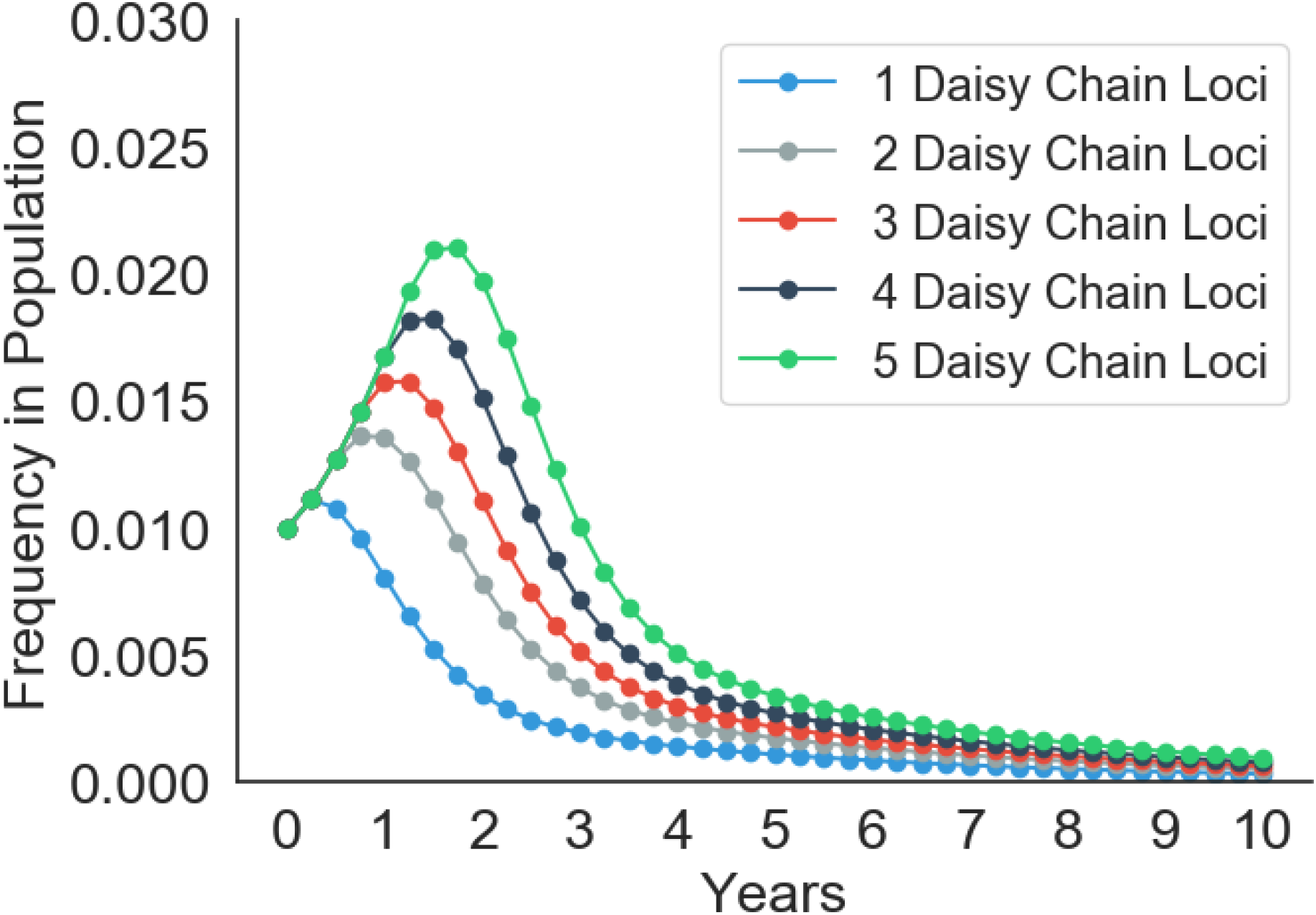
Forward simulations of daisy drive systems for the inheritance of GDMI designed with 1-5 daisy chain loci. Decay of the drive occurs after *n* generations, therefore more loci produce a longer lasting drive. However, because GDMI spreads slowly in the population compared to a fully outcrossed population, peak frequency of GDMI is low. Nearly 30 daisy chain loci are required to reach peak frequency of 50%, rendering daisy drive infeasible for implementation in this system.

### Dominance and penetrance of immune allele

The above simulation in Fig. S3 demonstrates a strong qualitative fit to empirical data from selection experiments. The frequency of infection, which is the fraction of infected snails out of the total surviving exposed individuals, is a phenotype resulting from immunity to a miracidial strain. The phenotype is a function of the exposure dose (number of miracidia) and the genetic underpinnings of immunity, including the number of loci involved, the dominance of immune alleles over susceptible alleles, and the penetrance of immune alleles. The genetic contributions to this phenotype could be myriad, but some large-effect loci have been identified in model snail species like *Biomphalaria glabrata*. These loci tend to be regions with high genetic variability and are linked to transmembrane proteins and receptors, which suggests a role in epitope recognition of an invading miracidium or sporocyst. One large-effect locus identified in Tennessen et al. 2015 served as a template for the default parameters used in the simulations in this work. We assumed a single locus model to represent this tightly linked gene cluster, and dominance of the immune allele over the susceptible allele was assumed complete as demonstrated in their empirical work. With a penetrance of 0.8, the model closely replicated observed evolution of immunity. However, as genetic work, such as genome wide association studies, identify new regions associated with immunity to schistosome infection in snails, more clarity will exist in the genetic contributions to immunity. Genes conferring immunity to one species or strain of schistosome may not confer immunity to others. These genes may not be conserved across snail species, and it is likely that immunity constitutes a wide array of variable genes. In *B. glabrata* two such polymorphic loci have been identified and described by Tennessen et al. 2015 and Tennessen et al. 2020 [57, 62]. Named Polymorphic Transmembrane Cluster 1 and 2 (PTC 1 and 2), these regions are each associated with several fold decreased odds of infection. However, the immune allele within PTC 2 likely has higher penetrance than the immune allele within PTC 1, and although the immune allele in PTC 1 is haplosufficient and completely dominant, incomplete dominance is observed for the immune allele within PTC 2. Variation in the genetic mechanisms of immunity can result in altered evolutionary trajectories in the face of the same strength of selection due to infectious miracidia. We show below how variation in dominance and penetrance changes the expected frequency of infection after generations of selection observed in Fig. S4.

**Figure 3:**
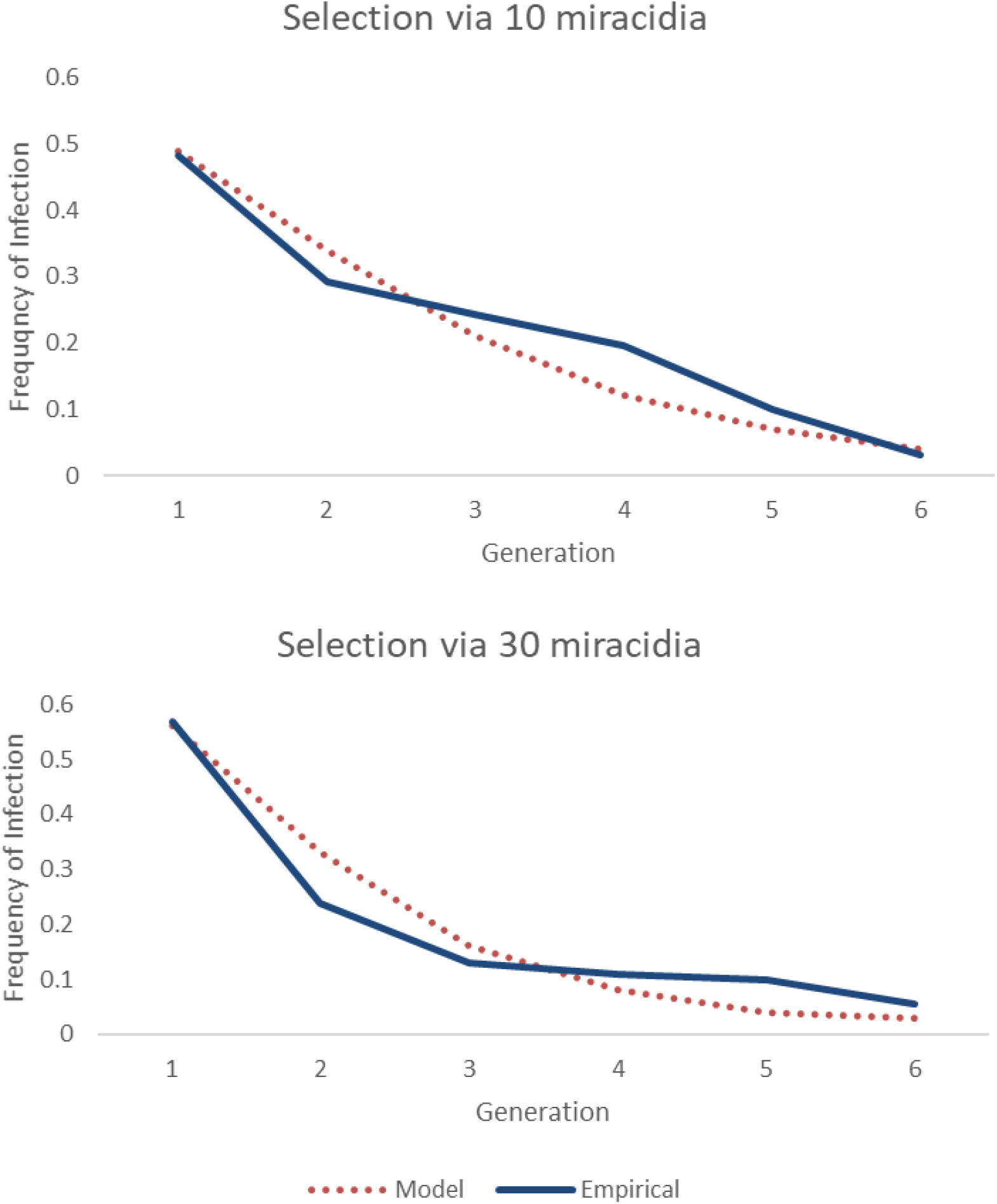
Qualitative comparison of model results to empirical data from published selection experiments by Tennessen et al. 2015. Two selection experiments were conducted, the first challenging each snail with 10 miracidia (top), and the second challenging each snail with 30 miracidia (bottom). Given initial conditions similar to experimental conditions, both models perform well in recapitulating selection for immunity to infection.

**Figure 4:**
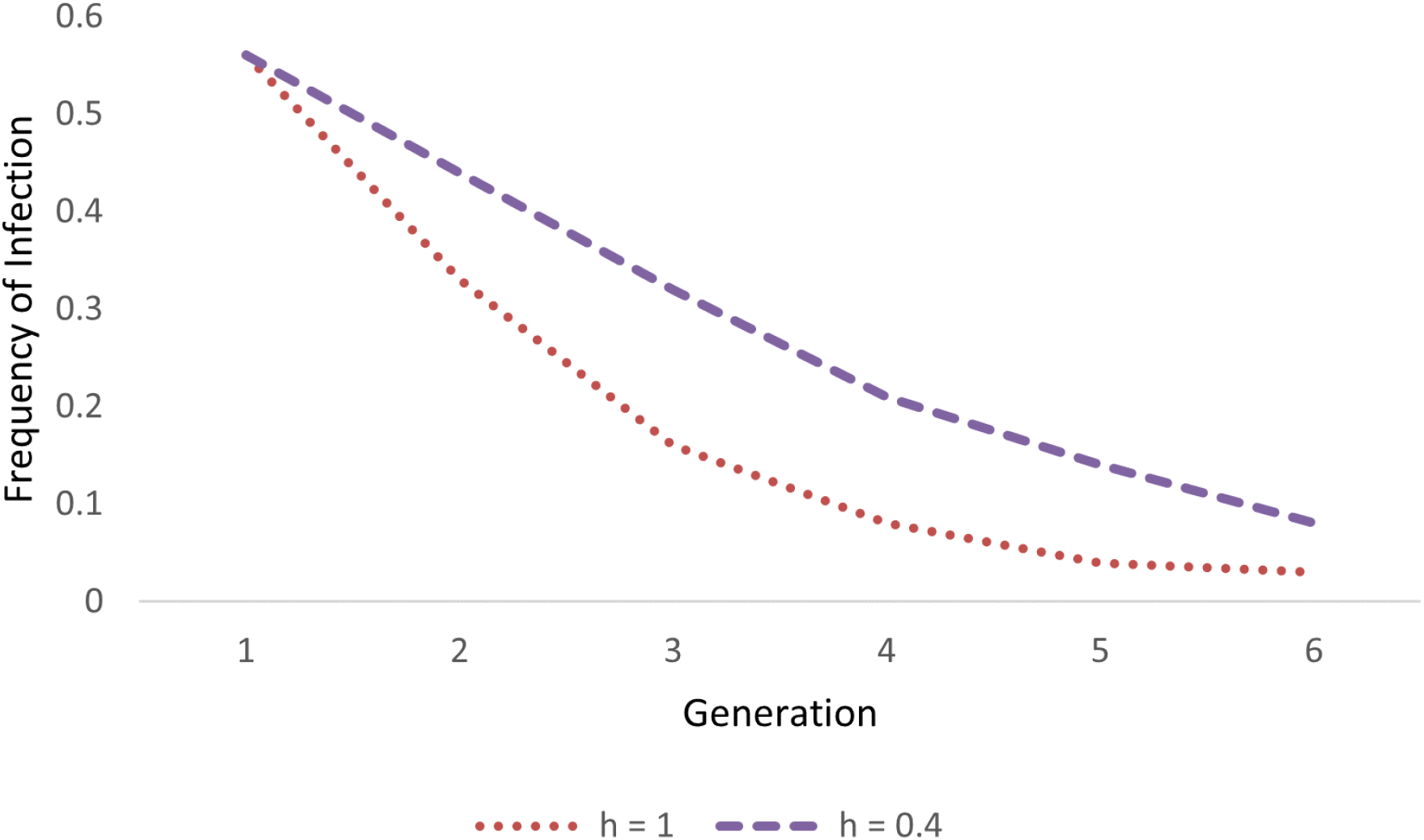
The relationship between the immune and susceptible alleles described by the dominance coefficient governs the trajectory of evolution for naturally-occurring immunity. Lower dominance of the immune allele leads to slower evolution of immunity, which could change the speed at which GDMI increases in frequency in a population.

In contrast to naturally-occurring alleles that whose inheritance is governed by the interaction between selection and dominance, an effective gene drive (high homing efficiency) may not be sensitive to this interaction because gene drive heterozygotes are produced at low frequencies, and therefore, dominance plays only a small role in determining the fitness of gene drive alleles. We simulate GDMI inheritance under default conditions to test whether GDMI frequency is significantly altered by the strength of dominance of the immune allele over the susceptible allele(s). Dominance coefficients of 1 and 0.4 were chosen as the measured upper and lower bounds for PTC 1 (h = 1) and PTC 2 (h = 0.4). The results in Fig. S5 show that the effect of dominance on the success of gene drive in the 10 year evaluation window is minimal. These results support the notion that an effective drive designed targeting either locus would operate similarly provided other factors are equivalent.

Two caveats may change these results: PTC 2 contains 2 susceptible alleles, therefore a natural heterozygote of both susceptible allele exhibits intermediate immunity compared to natural homozygotes of each susceptible allele, and penetrance of immunity associated with PTC 2 was measured higher than PTC 1 (approx. 2-fold higher odds ratio). In the case of two susceptible alleles displaying a range of immunity across natural susceptible genotypes, independent assortment ensures that relative fitness of immune alleles will depend on the average absolute fitness of the susceptible alleles. The average absolute fitness of susceptible alleles is a byproduct of their interactions to produce a range of susceptible phenotypes. One susceptible allele in PTC 2 is additive: the homozygote is twice as susceptible as the heterozygote (one susceptible allele, one immune allele). The other susceptible allele in PTC 2 is partially additive: the homozygote is less than twice as susceptible as the heterozygote. Barring other epistatic interactions, the measured susceptibility of the genotypes, their frequency, and the force of infection (directional selection) can be used to determine the relative fitness of immunity. However, differences between the alleles, including costs of maintaining each of the susceptible genotypes, is unknown and precludes investigation into differences between the fitness of the susceptible genotypes in the face of selection. This subject will require further empirical investigation to determine whether a spectrum of susceptible alleles may alter the speed of establishment of GDMI for a PTC 2 -like target. Based on results presented in Fig. 3 (main text), GDMI establishment is mildly sensitive to standing genetic susceptibility to infection, thus we expect minor differences in the evolutionary dynamics between a single susceptible allele system and a diversified susceptible allele system. A factor that has greater potential for impact on the evolutionary dynamics of GDMI is penetrance. Higher penetrance of immunity results in greater phenotypic variation with a heritable basis, which provides greater evolutionary potential. Immunity associated with PTC 1 is modeled with *ι* = 0.8. Susceptibility associated with PTC 2 represents up to 2-fold greater odds of infection. We simulate with *ι* = 0.9 in Fig. S6.

**Figure 5:**
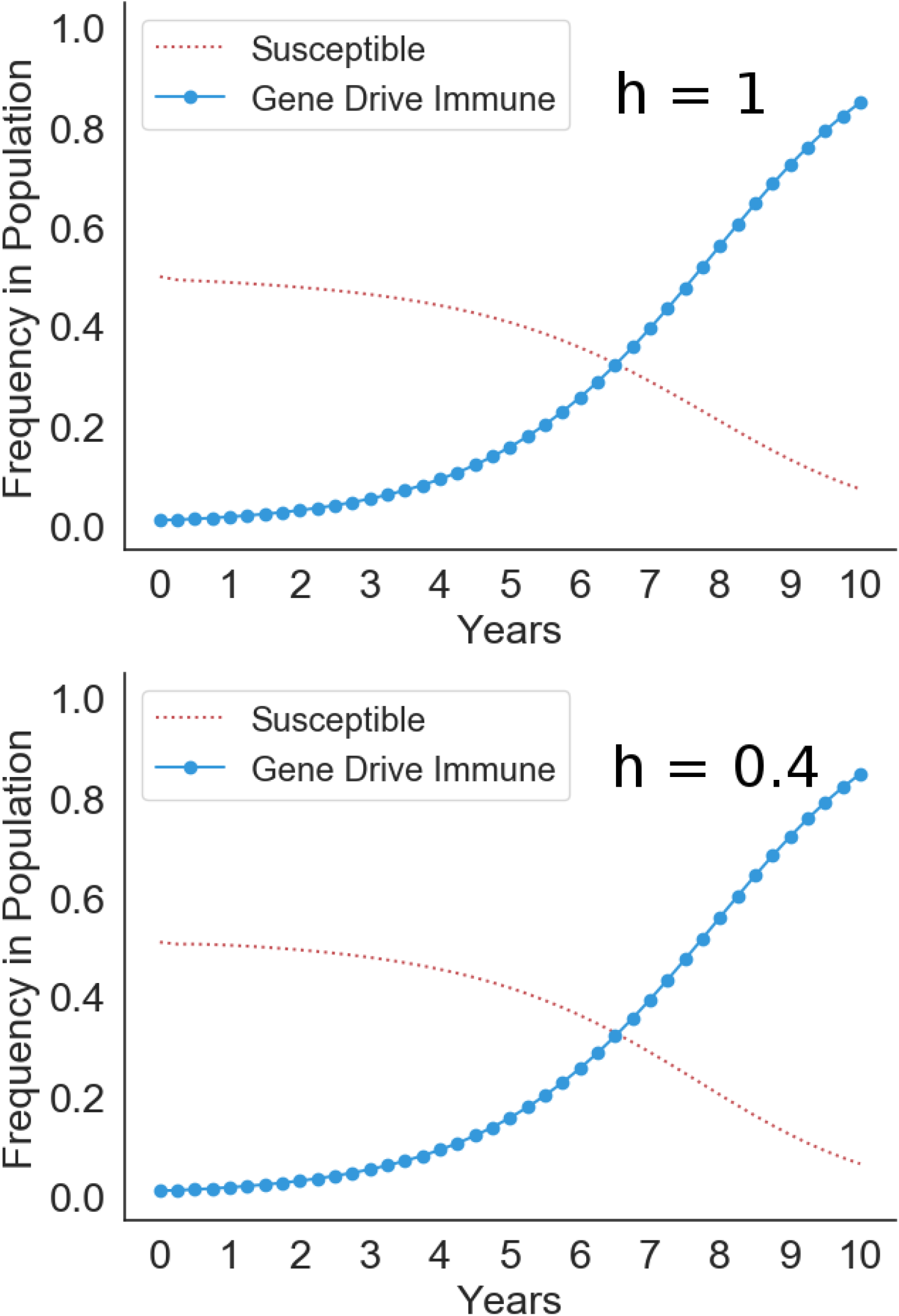
High dominance (top panel, h=1) representing PTC 1 and low dominance (bottom panel, h=0.4) representing PTC 2 do not yield measurably different results under default simulation conditions after 10 years.

**Figure 6:**
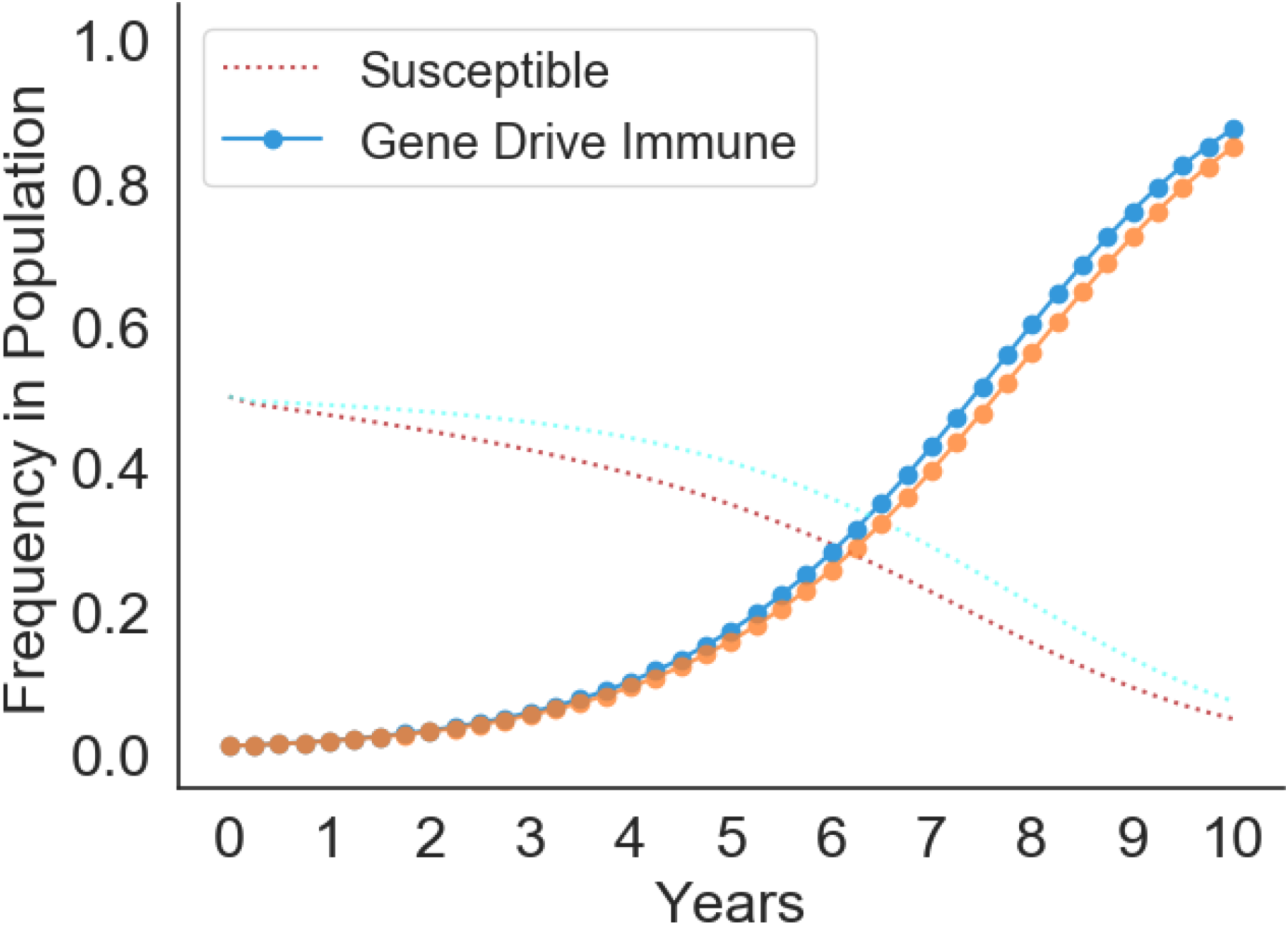
The effect of default penetrance (*ι* = 0.8) compared to higher penetrance (*ι* = 0.9) in the establishment of GDMI. Higher penetrance produces the blue GDMI and red susceptible lines, while lower penetrance produces the orange GDMI and light blue susceptible lines.

Despite the 2-fold greater odds of infection, PTC 2 susceptibility is not a significantly better target for GDMI. Resulting immunity in the population is similar after 10 years. These results change as the fitness advantages of GDMI over natural immunity diminish (e.g. low homing efficiency) and the fitness advantages of immunity over susceptibility strengthen (e.g. high force of infection).

### Generation time and population turnover

The evolutionary dynamics for GDMI reported here describe conditions in which the average time to reproduction is 3 months and the natural background mortality is half of the adult population in that time. However, when fortuitous environmental conditions prevail, or for snail species with shorter generation times, the establishment of GDMI in a population may happen at a different speed. In ten years, genotype frequencies in the population may be far different for these variable conditions. We demonstrate how variation of two basic life history parameters – natural mortality rate and generation time (mean time to reproduction) – influence the establishment of GDMI in 10 years. Fig. S7 displays simulations under default conditions, while these two life history parameters vary.

### Invasion conditions

Each genotype can be determined to be invading given that it is increasing in frequency at time *t* = 0. Invasion of a genotype does not guarantee increasing frequency at any time *t*, as conditions may change, even in the deterministic model (i.e. model results are not monotonic). However, invasion criteria are important determinants in understanding the behavior of the genetic system in the early stages of gene drive release or even in a natural but unstable genetic system (e.g. strong directional selection). For GDMI establishment, the relative fitness of the gene drive homozygote must be greater than 1. This can be directly determined by ensuring:

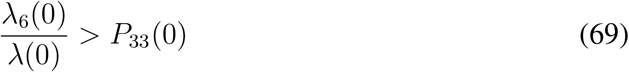

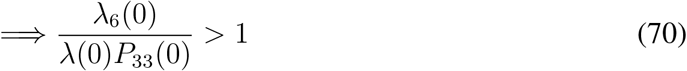

Additionally, the total population size must not decline towards extinction for this invasion to be successful. In Fig. S8 we calculate invasion thresholds for the variety of model parameters in relation to self-fertilization frequency.

**Figure 7:**
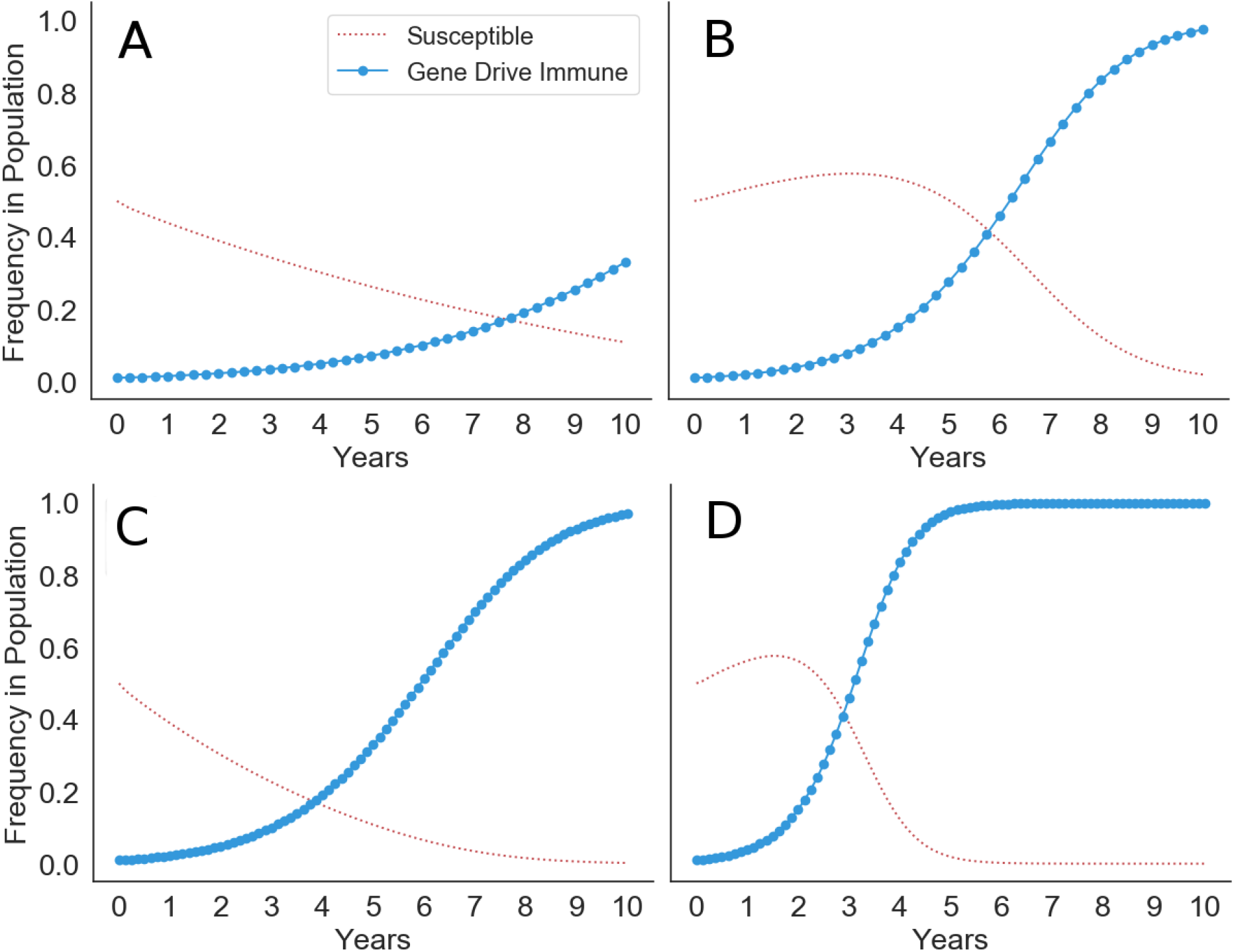
Simulations of susceptible and GDMI frequencies under variable life history strategies, namely mean generation time and death rate. Increasing death rate results in more population turnover each generation and more rapid fixation of GDMI. Panel A shows results for *µ* = 0.25 while panel B shows results for *µ* = 0.75. Similarly, shorter generations yields more rapid fixation of GDMI in 10 years because more generations occur within the time window. Panels C and D give show results for a mean generation time of 1.5 months (80 generations in 10 years) in contrast to 3 months (40 generations in 10 years). Panel C maintains *µ* = 0.25, and panel D maintains *µ* = 0.75.

**Figure 8:**
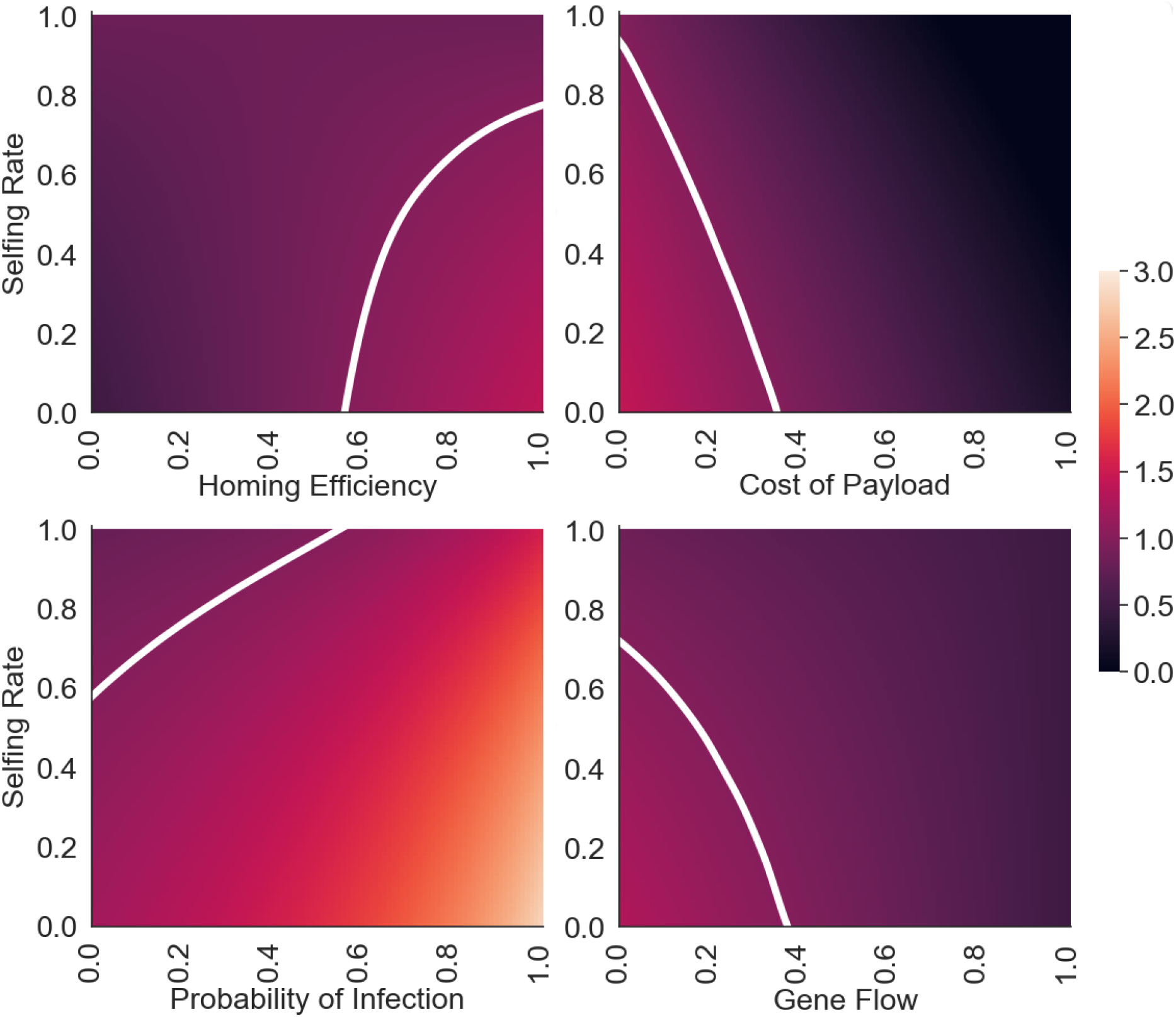
Invasion analyses for variables that influence the probability of invasion. Other parameters are held at their default value according to table S1, while the reproduction number is calculated as selfing rate varies. Lighter areas indicate higher reproduction numbers, and white lines represent the isocline at threshold conditions (*R*_0_ = 1). The ratio reported in equation S70 and *R*_0_ share a value of 1 under threshold conditions but are otherwise not precisely equal due to the nature of overlapping generations in the model.

Selfing rate and homing efficiency interact to form a curved region bounding values above *H* = 0.6 and below σ = 0.8. Homing efficiencies below 0.6 do not produce a viable gene drive under default conditions for any selfing rate. Similarly, selfing rates above 0.8 render a gene drive less fit than the natural population and unable to invade. Also supported by Fig. 3 in the main text, the cost of the payload is a strong determinant in the success of the invasion of GDMI. Perhaps strongest is the influence of gene flow on the invasion of GDMI, which restricts invasion to a small subset of conditions, rapidly excluding snail species with moderate selfing rates as gene flow increases. This does not capture long-term dynamics where GDMI individuals immigrate to the focus population – a process that may occur if GDMI establishes in the neighborhood of the focus population. Highlighted here is the robustness of invasion across the range of disease conditions. In both low and high transmission areas, invasion of GDMI is possible for some or all snail species. Species exhibiting high selfing rates may be invaded by GDMI in high transmission areas.

### Extinction risk

Invasion thresholds are valid and provide context for the conditions in which GDMI will proliferate given that the GDMI allele is not lost from the population due to genetic drift. Drift is strongest in generations immediately proceeding introduction (in a homogeneous environment) when the size of the pool of GDMI alleles is small. In contrast to a deterministic invasion process, genetic drift depends on the size of the seed GDMI population. We show how the probability of extinction of GDMI in 10 years (40 generations) varies with the size of the seed population and the absolute fitness of GDMI. We assume the probability of extinction is driven by a stochastic death process contributed through background mortality and infection prior to reproduction in each generation. Default parameters for background mortality and infection for GDMI homozygotes are 0.5 *gen^−^*^1^ and 0.03 *gen^−^*^1^, respectively. Therefore, we model the stochastic death process as *P r*(*X* = *k deaths*) = *B*(*n,* 0.53), where *n* is the number of GDMI homozygotes (excludes heterozygotes for simplicity due to their transiency at *H* = 0.9). Given a geometric mean absolute fitness *f* and number of generations from introduction, *t*, the cumulative distribution function representing the probability of extinction at time *t* is

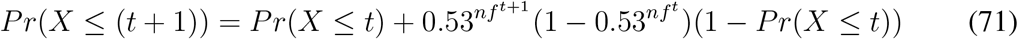

We calculate the probability of extinction within 40 generations (*t* = 40) using this recursive formulation and plot the results in Fig. S9 across a range of absolute fitness values and number of seeded GDMI individuals in the focal population.

Results indicate that the probability of extinction depends primarily on the absolute fitness of GDMI and little on the number of seeded individuals. Extinction is guaranteed below geometric mean absolute fitness of 0.9, regardless of the size of the introduced GDMI cohort. At low numbers, the threshold of extinction resides at a fitness of 1. This threshold is stark, with very little intermediate extinction risk in 40 generations. This is due primarily to the number of generations simulated; fewer generations would yield more intermediate extinction probabilities on the same plot. Combined with earlier results, this shows that the size of the introduced cohort is important for rapid fixation of GDMI but less important for persistence of GDMI in the population. This conclusion may not hold when reproduction and death is highly variable due to factors like seasonality. Higher variability will result in higher extinction risk, particularly for smaller seed populations.

### Seasonality

Dramatic variation in available snail habitat due to seasonal changes in precipitation is common in schistosomiasis endemic regions. Highest variation is observed in sites with ephemeral water bodies and agricultural areas. It is unclear how seasonal variation in habitat availability will alter the speed of establishment of GDMI. We compare GDMI establishment with and without seasonality in carrying capacity of the snail population, with a four fold change in carrying capacity simulated in the seasonally variable population. Fig. S10 demonstrates that seasonality slows the establishment of GDMI, even in a deterministic model. Although not shown in Fig. S10, higher variability in carrying capacity corresponds monotonically to slower establishment of GDMI. These results support the use of GDMI in sites with less seasonal variability in snail population abundance.

**Figure 9:**
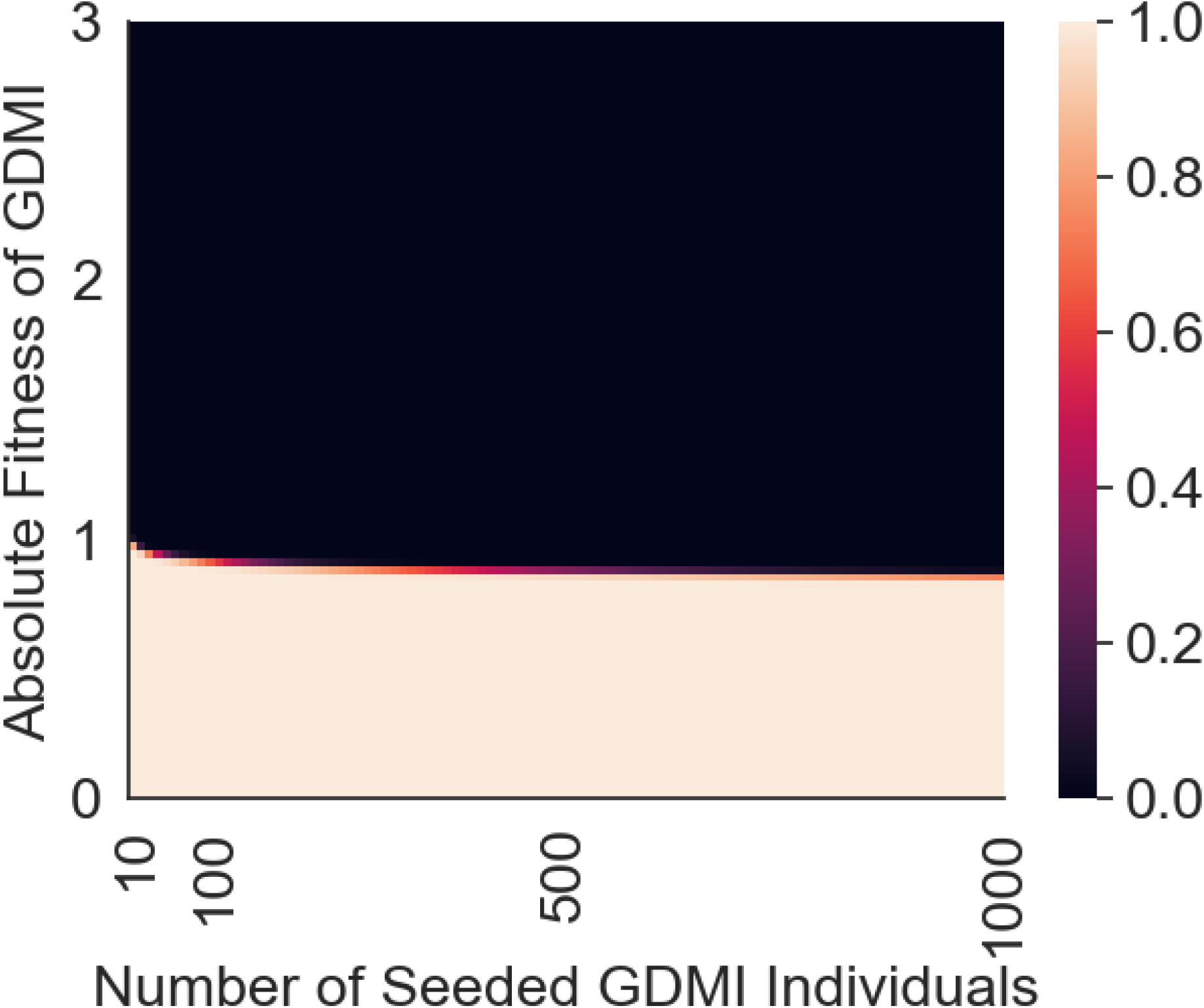
The probability of extinction within 40 generations according to absolute fitness and the number of seeded GDMI individuals. Darker values represent low likelihood of extinction.

**Figure 10:**
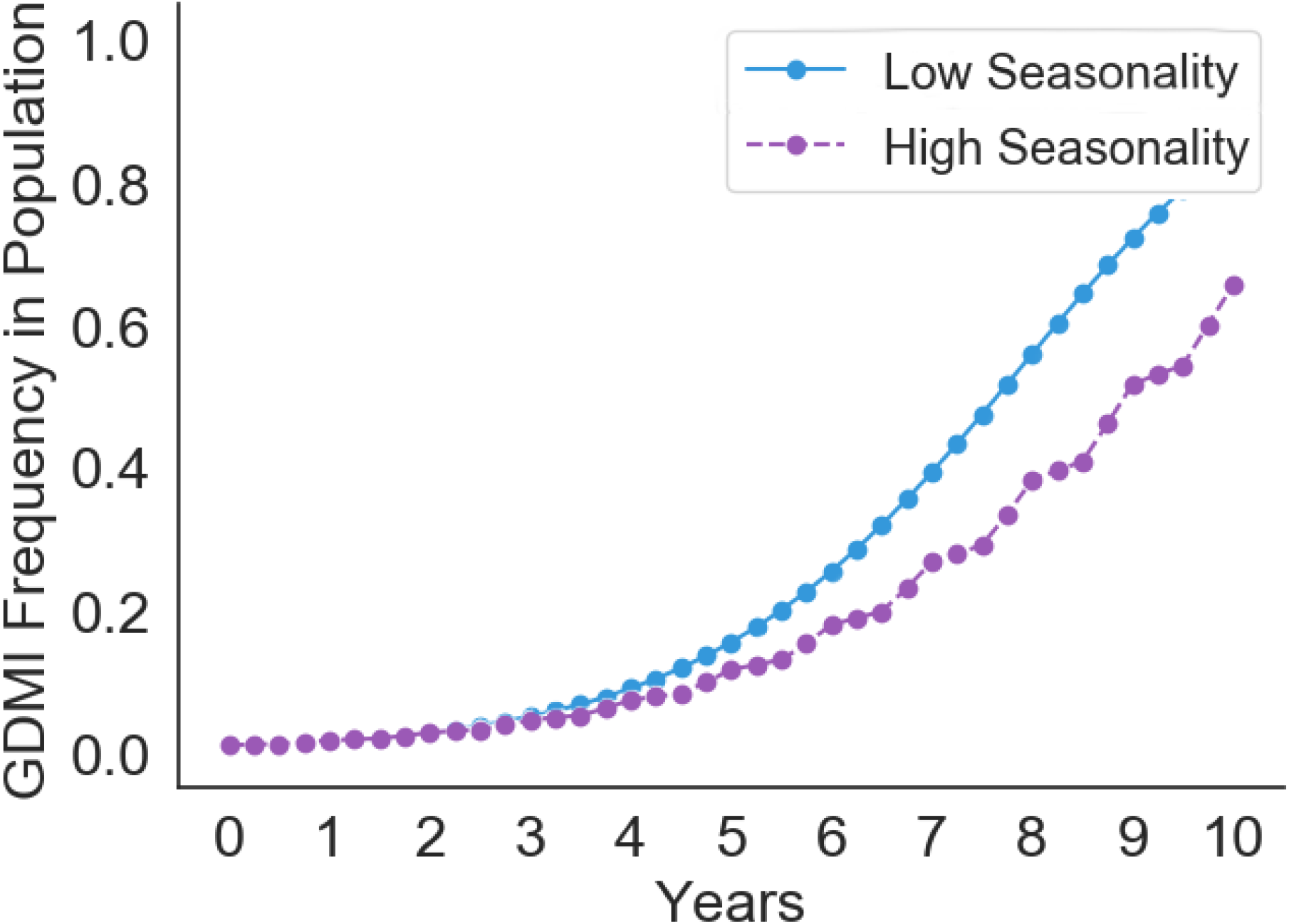
The spread of GDMI in a population with fluctuating carrying capacity due to seasonal rainfall and habitat variation. High seasonality assumes at 4 fold change in carrying capacity in 2 generations, with a full cycle occurring in 4 generations (equal to 1 year with default generation time): 200 %, 100 %, 50 %, 100% carrying capacity cycle. Low seasonality assumes no fluctuation in carrying capacity.

### Epidemiological model

Here we modify the classic MacDonald model for schistosome transmission to include the frequency of resistant snails that occurs in the environment due to introduction and subsequent spread of GDMI in the population. The simplest form this takes is to subtract the frequency of immune snails from the susceptible snail frequency such that the frequency of susceptible snails is given by (1 *− y − ρ*), where *ρ* is the frequency of immunity (natural and engineered). The resulting system of coupled ordinary differential equations is given below (equations 5 and 6 in the main text):

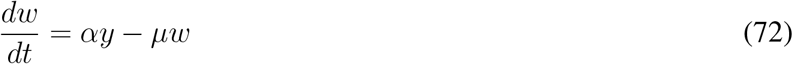

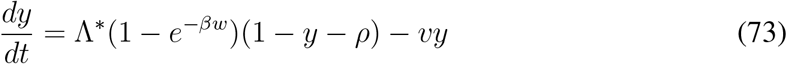

These two equations govern the prevalence of infection in snails, *y*, and the mean per capita worm burden in humans, *w*. The distribution of adult worms in the human population is assumed to approximate a negative binomial distribution. Parameters and their values are described in table S2.

We evaluate the efficacy of GDMI intervention by comparing mean worm burden after a ten year period with the mean worm burden at equilibrium endemic conditions. We calculate transmission rates Λ and *α* at endemic equilibrium. Let *w^*^, y^*^* be nontrivial equilibria for which 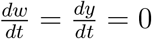.

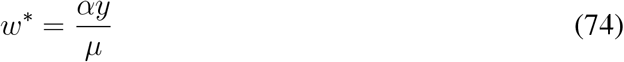

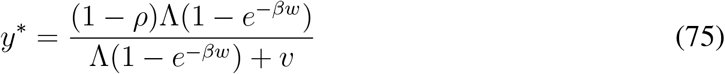

**Table 2:** Parameter values for the epidemiological model

| Parameter | Description | Value | Ref. |
| --- | --- | --- | --- |
| $\alpha$ | transmission rate converting snail infections to adult worms in humans | $142 \text{ wk}^{-1}$ | calculated based on model equilibrium at $w^* = 710$, $y^* = 0.02$ |
| $\mu$ | death rate of adult worms | $0.004 \text{ wk}^{-1}$ | Harmonic mean of range of $(3 - 7 \text{ yrs})^{-1}$ commonly reported in literature across *Schistosoma spp.* [63] |
| $\Lambda^*$ | force of infection from humans to snails at endemic equilibrium | $0.0104 \text{ wk}^{-1}$ | derived from epidemiological model |
| $\rho$ | fraction of immune snails | variable, $\rho^* = 0.5$ | determined by genetic model |
| $v$ | death rate of infected snails | $0.25 \text{ wk}^{-1}$ | Harmonic mean of death rates of infected *Bulinus globosus* and *Biomphalaria pfeifferi* [55] |
| $b$ | per capita worm to snail transmission rate | $3 * 10^{-5} w^{-1}$ | calculated based on model equilibrium at $w^* = 710$, $y^* = 0.02$ |
| $\beta$ | human population to snail transmission rate | variable, $\beta^* = 0.0147 \text{ wk}^{-1}$ | calculated: $\beta = \frac{1}{2}\phi b$ |
| $m$ | per capita mean number of mated pairs of adult worms | variable, $m^* = 348$ | calculated: $m = \frac{1}{2}\phi w$ |
| $k$ | clumping parameter: $NB(w, k)$ | 0.24 | fitted to *S. mansoni* data [64] |
| $w$ | per capita mean worm burden | variable, $w^* = 710$ | calculated based on model equilibrium at $\Omega = 0.80$ |
| $y$ | frequency of patent infections in snail population | variable, $y^* = 0.02$ | field-observed average in endemic regions [65] |
| $\phi$ | per adult worm mating probability | variable, $\phi^* = 0.98$ | calculation based on $NB(w, k)$ |
| $\Omega$ | per capita prevalence of at least one mated pair of adult worms | variable, $\Omega^* = 0.80$ | field-observed average in endemic and hyperendemic regions [66, 67] |

*y^*^* is frequently estimated in field surveys and may vary across sites or through the year due to seasonal variability in rainfall and human use of aquatic snail habitat like drainage areas, irrigation ditches, or natural water bodies. Despite variation, low infection prevalence (0-5%) in snails is observed, even in hyperendemic areas. Explanations for low prevalence are multifactorial and relate to the duration of patency, increased mortality rate of patent snails, heterogeneous exposure to miracidial infection, partially evolved immunity to infection, and competition with other trematodes. *w^*^* is not as easily estimated, as measurements rely on quantification of shed eggs in urine and fecal samples. The quantity of eggs is correlated but not linearly related to the number of paired worms, as human immunity leading to granulomatous formation around released eggs as well as potential interactions among adult worms and variability in egg production can obscure the relationship between eggs shed and worm burden. Human autopsies performed on known and suspected schistosomiasis cases reveal differential distribution of eggs and associated pathology with increasing intensity of infestation. Cheever (1968) observed that fewer eggs were present in the rectal mucosa and feces of *S. mansoni* infected individuals with associated fibrosis of the liver. This demonstrates that pathology, intensity, and egg count are not directly related, and the nature of their relationship requires biological knowledge of both the distribution of worms across tissue and the interactions between worms and the immune system. Despite these limitations, a reasonable heuristic is a 1:1 ratio of adult worm mated pairs and eggs per gram (EPG) in feces (*S. mansoni*). Multiple lines of evidence, including challenge experiments in mice, organ specific autopsies and perfusions, as well as observed distributions of EPG in human populations suggests that per capita mean worm burden (MWB) in highly endemic areas can exceed 1000. We simulate moderate-high endemicity with an infection prevalence of Ω*^*^* = 0.80. When the prevalence of infection is *<<* 1, a proportion of adult worms fail to pair with a mate and reproduce. The number of mated pairs can be calculated given the MWB and the distribution of adult worms in the human population. A negative binomial distribution is found to best represent the distribution of adult worms in humans. It is overdispersed, and dispersion increases as prevalence decreases. Prevalence of detectable eggs, and therefore successfully mated pairs is given as

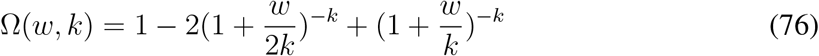

and *w* is calculated via substitution of the known prevalence of infection, Ω, and aggregation parameter, *k*, and solving numerically. An equal ratio of male and female schistosomes is assumed in this calculation, as is that the rates of transmission between the two sexes are equivalent [68]. We also assume that both sexes transmit together, and there is no sex-specific compartmentalization in the human body that would limit pairing of adult schistosomes. *w^*^* = 710 occurs at an endemic equilibrium prevalence, Ω, of 80%. Given *w^*^* = 710 and *y* = 0.02, *α* can be calculated as

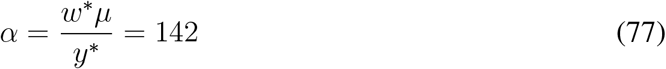

In contrast the *α*, which holds a constant value, *β* is a function of the distribution of worms in the local human population. The distribution changes non-linearly with worm burden and prevalence. For simplicity, we assume that *k* is invariant as worm burden and prevalence change, although evidence suggests higher aggregation with higher burden in some populations. *β* takes the form:

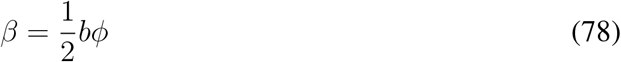

in which *b* is a transmission constant that relates the per capita number of mated worm pairs, *m*, to new infections in snails.

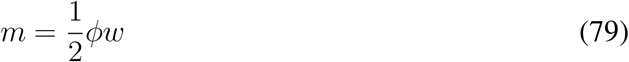

*φ* is the mating probability given by the negative binomial distribution where 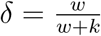.

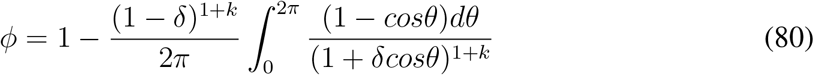

Given *w^*^* and *y^*^*, Λ*^*^* can be calculated by approximating that (1 *− e^−βw^*) *≈* 1 at endemic equilibrium conditions. Equation S73 simplifies to:

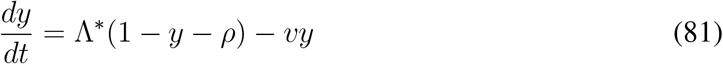

Solving for the nontrivial equilibrium yields:

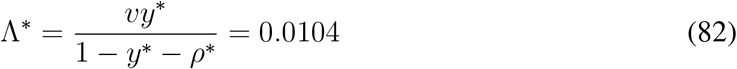

The probability of infection per generation, *P r*(*y*^+^), can be approximated from the force of infection and the differential equation for snail infection prevalence:

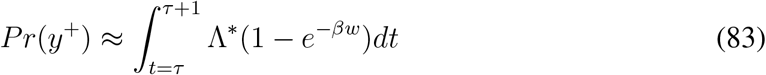

This expression represents the per capita number of snail infections expected in a susceptible population in a generation (*t* = *τ* weeks).

We do not yet have an estimate for *β* (variable) and therefore no estimate for *b* (constant). These values we determine by calibrating the model with known values for *R*_0_ in moderate-high transmission sites. The magnitude of *R*_0_ has never been precisely measured for schistosomiasis, as doing so would require measurements of innate immunity in the snail population. It is unknown whether genetic immunity provides cross protection for other trematode species, and therefore, even in a previously schistosome-naive area, pre-existing immunity requires measurement through challenge experiments. With this caveat in mind, *R*_0_ measurements are widely thought to exist in the range of 2-5 for schistosomiasis, likely exceeding 3 in moderate-high transmission sites [69]. In a fully susceptible snail population, these values will be higher, and in all likelihood empirically measured *R*_0_ values underestimate true *R*_0_ values predicated on a fully susceptible host population. From our system of differential equations we calculate the effective reproductive number *R_t_* and from it, derive the *R*_0_ under conditions of partial immunity in snails to calibrate *β*. Linearizing the system of equations with respect to *w* and *y*, we form the transmission and transition matrices outlined by Diekmann et al. in their next generation matrix (NGM) approach to calculate *R*_0_ [70]. We extend this approach by relaxing the assumption that the populations of snails and humans are fully susceptible and that no disease is present before an index case. Doing so, we calculate transmission and transition matrices for *R_t_* as:

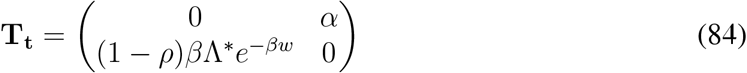

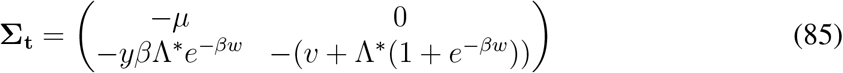

We calculate the time-varying NGM as:

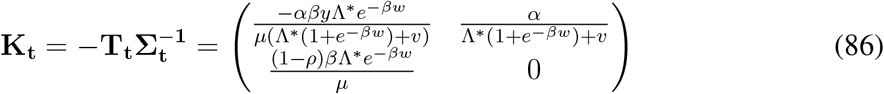

The expression for *R_t_*, computed as the spectral radius of **K_t_**, is

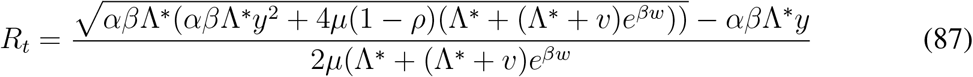

A derivation of *R*_0_ would require setting *ρ* = *w* = *y* = 0. However, because empirical measurements of *R*_0_ have not accounted for variations in *ρ*, we calibrate *β* from an empirical form of this equation. Specifically, we set *ρ* = 0.5 according to default conditions that are based on empirical measurements of innate immunity in field captured snails. Setting *w* = *y* = 0 yields the following expression for an empirical *R*_0_

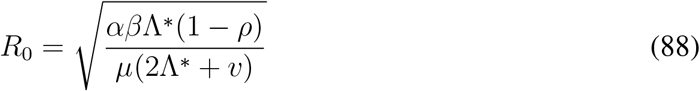

and when *ρ* = 0.5, the expression becomes:

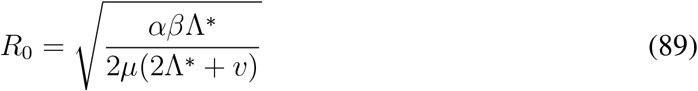

Solving for *β* yields

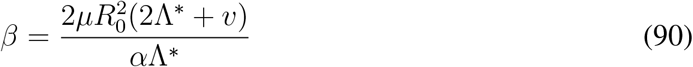

Recall that *β* is a function of the negative binomial distribution of worms, which determines the probability of mating success among adult worms. However, this theoretical construct breaks down for the low numbers assumed in an index case. For at low numbers, *φ* would approximate zero for a stationary *k* = 0.24, and extinction is predicted. The concept of *R*_0_ would be irrelevant for schistosomiasis if these theoretical predictions were valid. Instead we assume that early transmission of cercariae are highly clustered and that *φ* remains high for the purposes of estimating the constant *b* which scales *β*. Setting *φ* = 1, we achieve

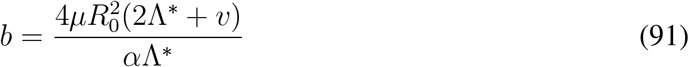

We set *b* to a value with one significant digit so that *R*_0_ is approximately the median of empirically measured values. This gives *b* = 0.03 and *R*_0_ = 3.2. In practical terms, *b* represents the ‘rebound speed’, which is the pace infections can accrue after chemotherapy treatment. Under certain conditions, our estimate may represent the low end of this rebound speed due to the assumption of 100% mating success in index case infections. Moreover, our estimate of *R*_0_ = 3.2 is calibrated based on prior empirical measurements, which almost certainly underestimate the true *R*_0_ as specified by a fully susceptible host population. We do not explicitly account for adaptive immunity in humans, which has been shown to increase over 4410+ years into adulthood. Accounting for evolved innate immunity in snails by setting *ρ* = 0, we find that *R*_0_ = 4.5.

In Fig. S11 we show the long-term behavior of the default model without introduction of GDMI. A reproduction number of 3.2 produces a rapid rise of an epidemic past the endemic equilibrium, and as the snail population evolves immunity, an equilibrium is established. Feedback from schistosome transmission produces stabilizing selection on immunity in snails.

In Fig. 4 of the main text, GDMI was evaluated in comparison to and with coincident annual MDA treatment. 60% reduction in MWB in the population was modeled for each treatment and is a product of coverage and efficacy of the chemotherapy. Although alone GDMI is not capable of eliminating schistomiasis locally within a 10 year evaluation period, it was shown to successfully complement MDA under simulated conditions to produce greater and more sustained reduction than MDA alone. However, these results may be sensitive to several factors, especially the force of infection to humans which determines how rapidly the human population becomes infected from an infected snail population. Rapid reinfection results in a faster rebound to pre-treatment MWB, and therefore, subsequent treatment is less effective because MWB reduction is not long lasting. We explore high and low transmission conditions by manipulating *b*, which in turn, changes *β*. Fig. S12 shows the difference between *R*_0_ = 2.3 and *R*_0_ = 4.5 conditions as MDA and GDMI are applied.

GDMI performs favorably in absolute reduction in joint use with MDA when transmission is higher, indicating that the efficacy of GDMI is enhanced in conditions that are challenging for reduction through MDA alone. Additionally, the intensity of MDA treatment may have a strong effect on the benefits of GDMI, as selection pressures are changed. Fig. S13 displays the difference in reduction of MWB between low and high intensity MDA use under equivalent GDMI application.

**Figure 11:**
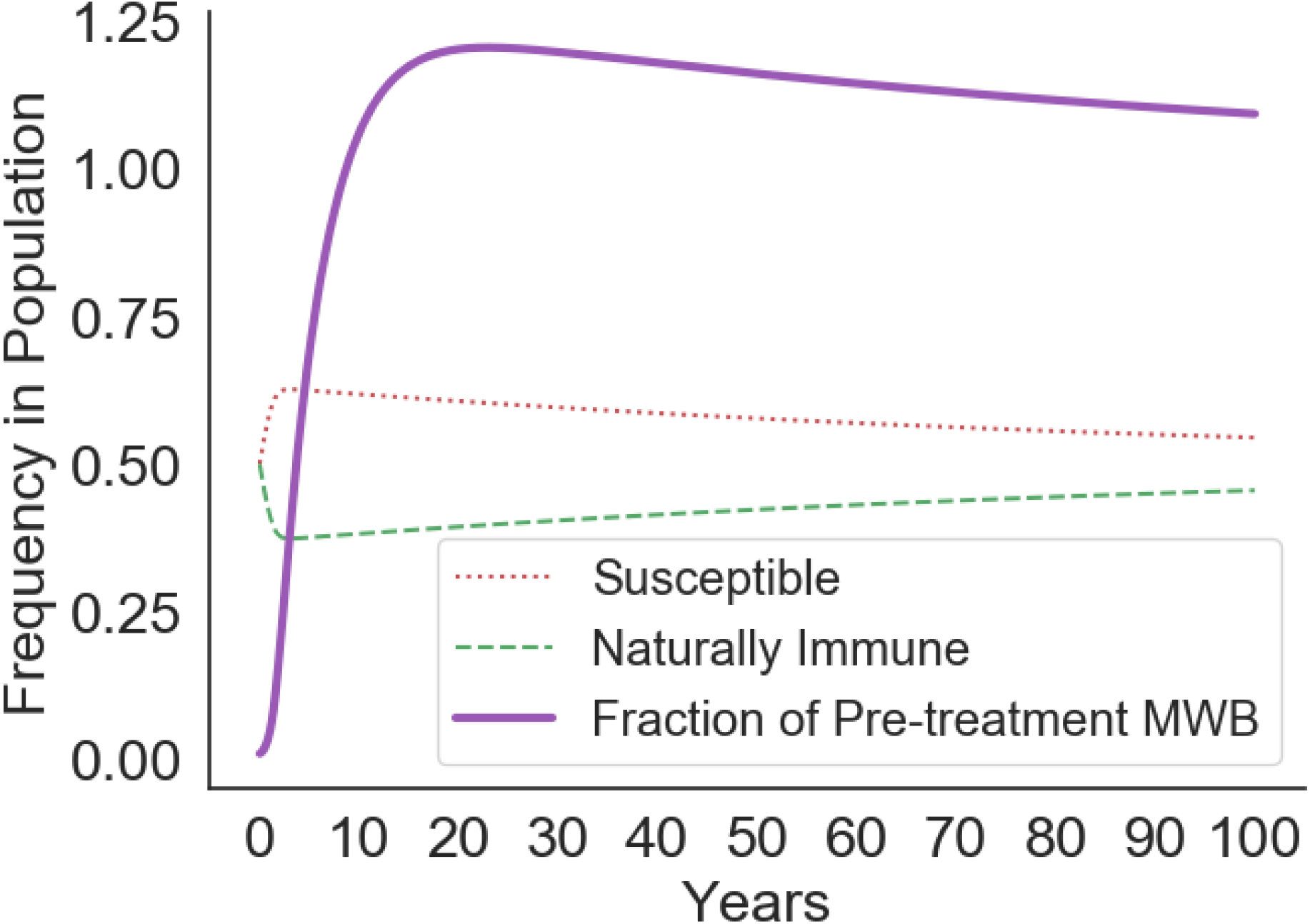
Simulation of the emergence of a schistosomiasis epidemic under default conditions. GDMI is not present, and long-term behavior of the model is observed to overshoot endemic equilibrium conditions and return to equilibrium over the course of many years. Susceptibility in snails is advantageous at low levels of infection early in the epidemic and is disadvantageous above equilibrium conditions.

**Figure 12:**
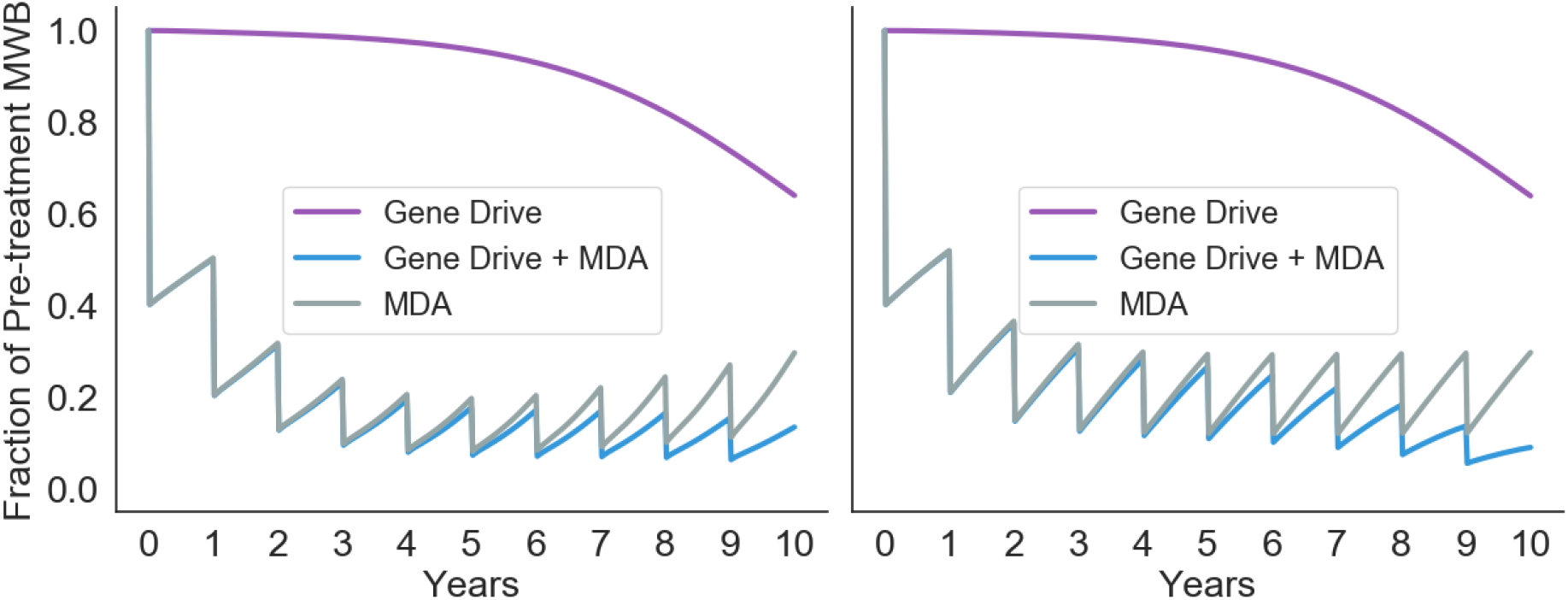
Comparative results among three treatment regimes under high and low transmission conditions. *b* is half of default conditions (left) and *R*_0_ = 2.3, producing slower rebounds after annual MDA treatment. More rapid rebounds are observed when *b* is twice default conditions and *R*_0_ = 4.5 (right).

These results demonstrate diminishing returns for the application of MDA at higher concentrations as immunity in snails evolves to favor higher transmission conditions when adult worms are eliminated quickly. Success of GDMI is slowed when force of infection on snails, and therefore positive selection on immunity, is reduced.

The treatment window of 10 years is common for evaluating funded public health campaigns, though results of this study will differ using longer treatment windows. We extend this window to 40 years to demonstrate the long-term effects of each of the treatment regimes. Additionally, we show that when MDA is remitted after 10 years, GDMI is able to maintain reductions in MWB, while without GDMI MWB returns to endemic equilibrium conditions (after an overshoot also depicted in Fig. S11).

**Figure 13:**
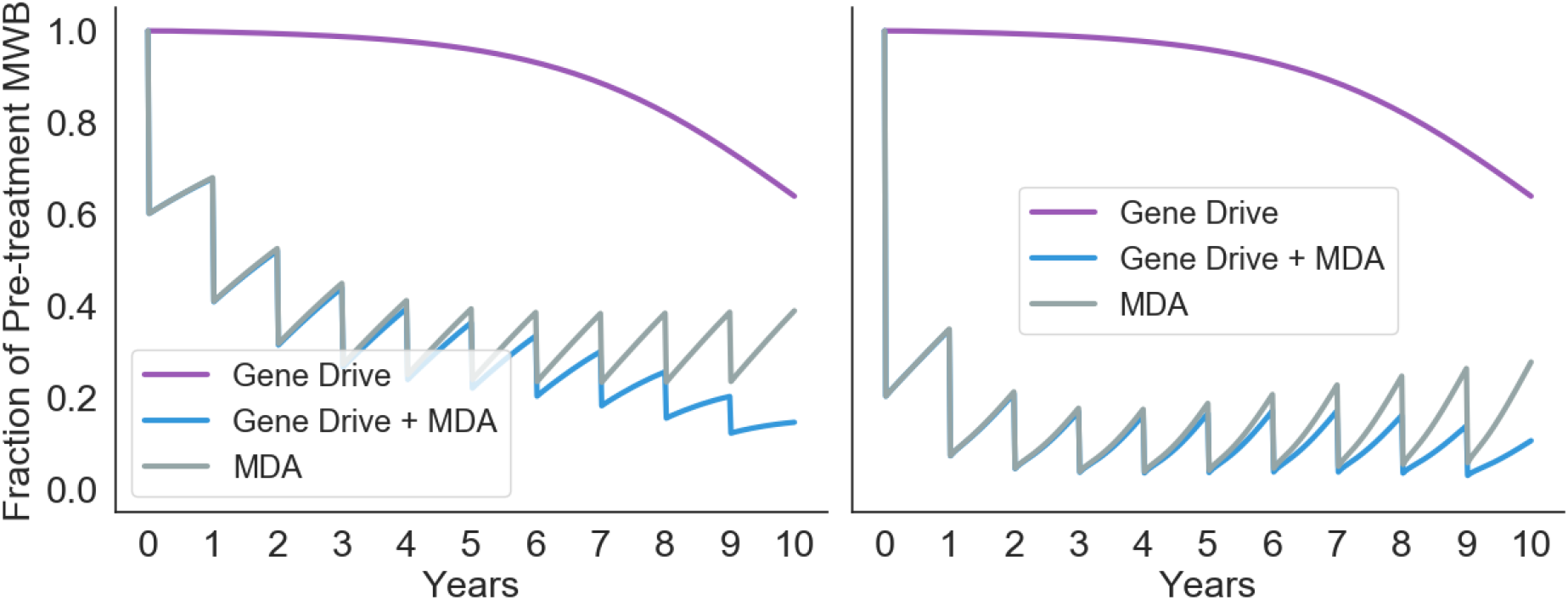
Comparative results among three treatment regimes under high and low intensity MDA application in the human population. 40% annual reduction in MWB (left) produces slower elimination across all treatment regimes compared to 80% annual reduction (right). Rebounds are concave down and relatively smaller for lower intensity MDA and concave up for high intensity MDA. This reflects slower loss of immunity, and for joint treatment the faster gain of GDMI, in the snail population due to higher selection pressure in favor of immunity in higher transmission conditions.

**Figure 14:**
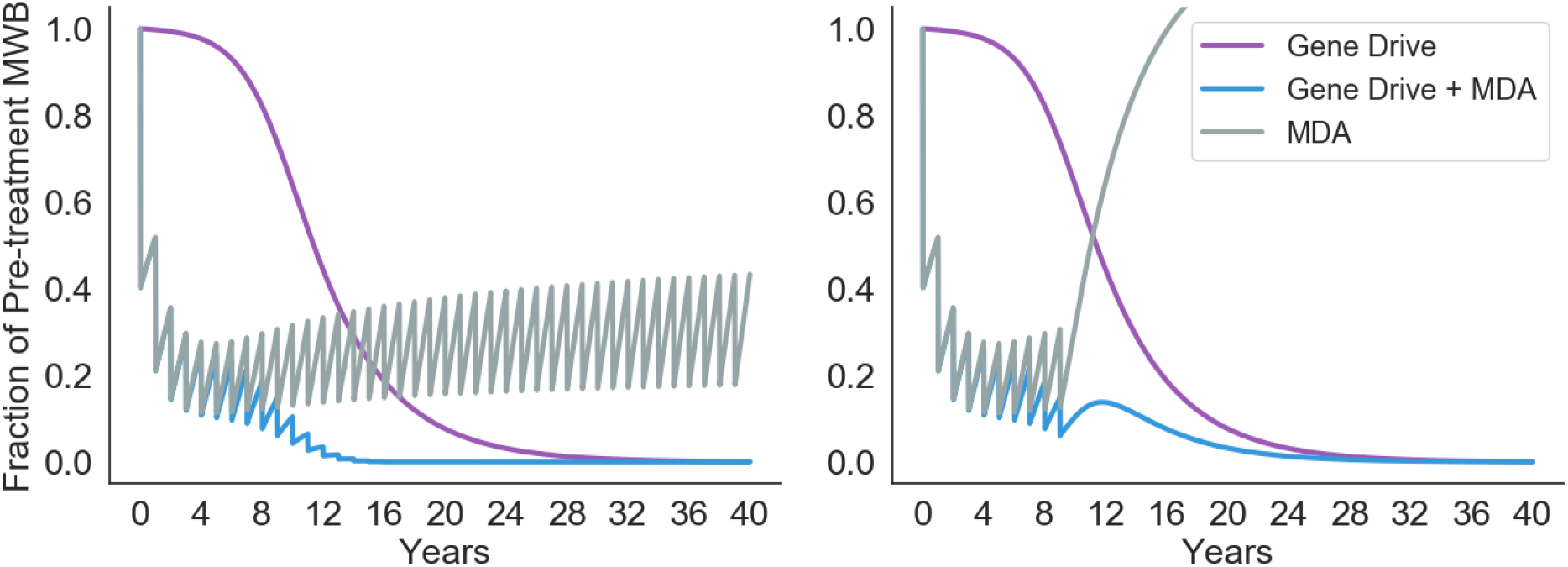
Simulations of the three treatment regimes for 40 years. MDA is continued annually for the duration of the simulation (left). MDA is stopped after 10 years of treatment (right).

